# Sex differences in deep brain shape and asymmetry persist across schizophrenia and healthy individuals: A meta-analysis from the ENIGMA-Schizophrenia Working Group

**DOI:** 10.1101/2024.10.24.619733

**Authors:** Delaina B. Cimmino, Brinley Zabriskie, Steven Luke, Boris Gutman, Dmitry Isaev, Kathryn Alpert, David Glahn, Amanda Rodrigue, Sinead Kelly, Godfrey Pearlson, Vince Calhoun, Stefan Ehrlich, Ole Andreassen, Diana Tordesillas-Gutierrez, Benedicto Crespo-Facorro, Theodore Satterthwaite, Raquel Gur, Ruben Gur, Gianfranco Spalletta, Fabrizio Piras, Gary Donohoe, Colm McDonald, Edith Pomarol-Clotet, Raymond Salvador, Andriana Karuk, Aristotle Voineskos, Peter Kochunov, Stefan Borgwardt, Ingrid Agartz, Erik Jonsson, Tilo Kircher, Frederike Stein, Katharina Brosch, Igor Nenadic, Felice Iasevoli, Giuseppe Pontillo, Andrea de Bartolomeis, Annarita Barone, Maraiateresa Ciccarelli, Annabella Di Giorgio, Arturo Brunetti, Sirio Cocozza, Mario Tranfa, Anthony James, Mojtaba Zarei, Morgan Hough, Lena Flyckt, Geraldo F. Busatto, Pedro G. P. Rosa, Mauricio H. Serpa, Marcus V. Zanetti, Theo van Erp, Adrian Preda, Dana Nguyen, Paul Thompson, Jessica Turner, Lei Wang, Derin Cobia

**Author notes:** Corresponding Author: Derin Cobia, PhD, Department of Psychology and Neuroscience Center, Brigham Young University, 1036 KMBL, Provo, UT 84602.

## Abstract

**Background:** Schizophrenia (SCZ) is characterized by a disconnect from reality that manifests as various clinical and cognitive symptoms, and persistent neurobiological abnormalities. Sex-related differences in clinical presentation imply separate brain substrates. The present study characterized deep brain morphology using shape features to understand the independent effects of diagnosis and sex on the brain, and to determine whether the neurobiology of schizophrenia varies as a function of sex.

**Methods:** This study analyzed multi-site archival data from 1,871 male (M) and 955 female (F) participants with SCZ, and 2,158 male and 1,877 female healthy controls (CON) from twenty-three cross-sectional samples from the ENIGMA Schizophrenia Workgroup. Harmonized shape analysis protocols were applied to each site’s data for seven deep brain regions obtained from T1-weighted structural MRI scans. Effect sizes were calculated for the following main contrasts: 1) Sex effects;2) Diagnosis-by-Sex interaction; 3) within sex tests of diagnosis; 4) within diagnosis tests of sex differences. Meta-regression models between brain structure and clinical variables were also computed separately in men and women with schizophrenia.

**Results:** Mass univariate meta-analyses revealed more concave-than-convex shape differences in all regions for women relative to men, across diagnostic groups (*d* = -0.35 to 0.20, SE = 0.02 to 0.07); there were no significant diagnosis-by-sex interaction effects. Within men and women separately, we identified more-concave-than-convex shape differences for the hippocampus, amygdala, accumbens, and thalamus, with more-convex-than-concave differences in the putamen and pallidum in SCZ (*d* = -0.30 to 0.30, SE = 0.03 to 0.10). Within CON and SZ separately, we found more-concave-than-convex shape differences in the thalamus, pallidum, putamen, and amygdala among females compared to males, with mixed findings in the hippocampus and caudate (*d* = -0.30 to 0.20, SE = 0.03 to 0.09). Meta-regression models revealed similarly small, but significant relationships, with medication and positive symptoms in both SCZ-M and SCZ-F.

**Conclusions:** Sex-specific variation is an overriding feature of deep brain shape regardless of disease status, underscoring persistent patterns of sex differences observed both within and across diagnostic categories, and highlighting the importance of including it as a critical variable in studies of neurobiology. Future work should continue to explore these dimensions independently to determine whether these patterns of brain morphology extend to other aspects of neurobiology in schizophrenia, potentially uncovering broader implications for diagnosis and treatment.

**Key Points:**

1. Statistical analyses revealed significant main effects for diagnosis and sex in deep brain shape morphology. Among patients with schizophrenia, there was a pattern of thinning and surface contraction in the bilateral hippocampus, amygdala, accumbens, and thalamus, and a pattern of significant thickening and surface expansion in the bilateral putamen and pallidum compared to healthy control participants. Between males and females, there was a pattern of significant thinning and surface contraction in the bilateral thalamus, pallidum, putamen, and amygdala in females compared to males.
2. There was no significant interaction between diagnosis and biological sex, suggesting that sex differences in deep brain shape and asymmetry among patients with schizophrenia reflect those observed in healthy individuals.
3. Small but statistically significant relationships exist between brain structure and clinical correlates of schizophrenia were similar for both men and women with the disease, such that higher CPZ was associated with shape-derived thinning and surface contraction in the caudate, accumbens, hippocampus, amygdala, and thalamus, and elevated positive symptoms were associated with shape-derived thinning and surface contraction in the bilateral caudate, right hippocampus, and right amygdala.

## Introduction

Schizophrenia is characterized by a disconnect from reality that manifests in the presence of positive, negative, and cognitive symptoms which vary in intensity across the lifespan and cause significant impairment in multiple functional domains (Craighead et al., 2017). Neurobiological abnormalities have been observed as a key feature of schizophrenia including cortical, deep brain, and cerebellar regions (Egloff et al., 2018). The extent to which structural changes in deep brain regions map on to the clinical presentation of the disorder is still largely unclear. Key deep brain nuclei that are consistently implicated in the pathophysiology of schizophrenia include the hippocampus, thalamus, hypothalamus, amygdala, basal ganglia, cingulate gyrus, and cerebellum, which play an important role in behavioral and emotional responses (Gutman et al., 2021). Overall, this presentation suggests a network-based neurobiological contribution to the etiology of psychosis given the inter- and intra-connected nature of these regions. Current research efforts continue to study how structural changes to various cortical-subcortical circuits influence the onset of the disorder, progress over time, and change the presentation of the disorder, which fluctuates in intensity and continues over the lifespan.

### Sex Differences in Schizophrenia: Clinical Presentation

Previous work highlights how schizophrenia presents differently in men and women. Age of onset is typically earlier in men, who may begin experiencing symptoms between 18 and 25 years old, compared to women, who demonstrate functionally impairing symptoms closer to 30 years old or even after 45 years of age (Li et al., 2016; Ochoa et al., 2012). As premorbid functioning is an important predictor of prognosis, it is important to note that women tend to exhibit higher premorbid adjustment and social support (Giordano et al., 2021; Ochoa et al., 2012). A more severe pattern of negative symptoms is generally observed in men, including blunted affect, avolition, and anhedonia, whereas women tend to present with increased mood disturbance and elevated manic symptoms (Giordano et al., 2021; Li et al., 2016; Irving et al., 2021). Among those who are actively taking antipsychotics, female patients tend to show better compliance and better outcomes with pharmacological interventions for psychosis, although they may also be at greater risk for side effects compared to male patients (Li et al., 2016). These differences may be mediated by the amount of drug entering the brain, as women require, on average, a lower dosage to achieve the desired effects (Seeman, 2021).

### Sex Differences in Schizophrenia: Neuroanatomical Findings

Recent studies have also examined sex differences in brain abnormalities observed in schizophrenia, suggesting a reconceptualization of the disorder that may explain differences in clinical presentation between men and women with the disease (Guma et al., 2017; Gutman et al., 2021; Egloff et al., 2018). Lateralization of dysconnectivity appears to be a focus of current research, finding that male patients tend to show a pattern of connectivity deficits that are left-lateralized and highlighted in striato-cortical systems, while the deficits in female patients are primarily right-lateralized (Wang et al., 2019). In terms of structural abnormalities, differences in shape deformation have been observed in the hippocampus and amygdala between men and women with schizophrenia (Egloff et al., 2018), although these findings appear to be inconclusive as other studies have found no significant sex differences in the brain (Guma et al., 2017). Furthermore, both men and women with schizophrenia demonstrate limbic asymmetry, including the hippocampus, amygdala, and thalamus, compared to individuals without the disorder, highlighting the emotional and behavioral impairments that arise due to the limbic system’s basic survival functions in the brain (Gutman et al., 2021).

Researchers have long hypothesized the relationship between brain abnormalities and clinical symptoms in schizophrenia, and that observed differences in clinical presentation between men and women suggests potentially unique sex-specific neurobiology (Guma et al., 2017; Li et al., 2016; Lang et al., 2018). Across men and women who present with psychosis, prominent biomarkers included enlarged ventricles and a reduction in hippocampal volume (Salminen et al., 2021). However, larger ventricles and smaller frontal and temporal volumes have been observed in males with schizophrenia, whereas females have exhibited smaller volumes of the anterior cingulate cortex and insula (Turkozer et al., 2020; Salminen et al., 2021). Furthermore, Guma and colleagues (2017) found an association between the intensity of negative symptoms observed in males with schizophrenia and structural changes in the amygdala. There is also some indication that women with schizophrenia experience greater white matter pathology relative to healthy women compared to schizophrenia versus healthy men (Kelly et al., 2018). While sex-specific variation in the clinical presentation of psychosis has received attention in the literature, a comparative examination of the underlying mechanisms contributing to these findings are unclear and warrant further research.

### Sex Differences In Healthy Individuals

Sex is biological trait known to significantly contribute meaningful variability in brain structure, which has been highlighted in decades of neuroimaging literature. Most frequently observed is larger intracranial (ICV) in men relative to women, which partially contributes to regional sex differences in brain volume (Jahanshad & Thompson, 2017) and persists throughout development (Kaczkurkin, Raznahan, & Satterthwaite, 2019). In a meta-analysis by Ruigrok and colleagues (2014) increased volume in bilateral amygdala, hippocampus, and putamen was observed in males relative to females, but with larger thalamus in females. Recent work has also identified diffusion-related microstructural sex differences in deep brain regions, including the hippocampus, thalamus and nucleus accumbens (Pecheva et al., 2024). Given the consistency of sex-specific variability in brain structure, and its potential import as an imaging phenotype for understanding psychiatric disorders (Salminen et al., 2022), a focused investigation of these features in schizophrenia could aid in determining whether deep brain circuitry contributes to the unique clinical presentations in men and women with schizophrenia.

### Aims and Hypotheses

The present study sought to systematically investigate sex differences, and related clinical correlates, in deep brain structures implicated in the pathophysiology of schizophrenia using high-dimensional brain mapping procedures. Shape deformation of deep brain regions was quantified using surface metrics in men and women with and without schizophrenia from multiple worldwide datasets. Using a meta-analytic approach, the first goal of the study was to examine the main effects of sex and diagnosis across four groups: men with schizophrenia, women with schizophrenia, healthy men, and healthy women. Previous work from our group (Gutman et al., 2022) revealed that patients with schizophrenia demonstrated abnormal shape deformation in the amygdala, hippocampus, thalamus, and basal ganglia (i.e., caudate, putamen, globus pallidus, and accumbens) compared to healthy control participants. For the next step, it was hypothesized that females would demonstrate greater concave shape in the hippocampus, amygdala, and putamen, but greater convex in the thalamus, compared to males.

The second goal of the study was to examine whether the interaction between sex and diagnosis had meaningful effects on the pattern of deep brain morphology in these samples. Given the differences in clinical presentation between men and women with schizophrenia (Giordano et al., 2021; Li et al., 2016), it was hypothesized that the effect of diagnosis on deep brain shape would be dependent on biological sex. In other words, it was anticipated that males and females with schizophrenia would exhibit different patterns of shape deformation in deep brain nuclei that were unlike sex differences observed among healthy individuals. Specifically, deep brain morphology in the amygdala, hippocampus, thalamus, and basal ganglia would vary among males and females with schizophrenia, such that greater deformation of the caudate and hippocampus would be observed in males, with greater deformation of the amygdala in females. Furthermore, given the progression of schizophrenia has also been associated with abnormal patterns of asymmetry (Okada et al., 2016; Schijven et al., 2023), how asymmetry across deep brain regions varies as a function of sex was also examined.

The third goal of the study was to evaluate sex-specific relationships between shape deformation of these deep brain regions and clinical dimensions of the illness in participants with schizophrenia. Namely, self-reported severity of positive and negative symptoms to determine whether unique shape abnormalities relate to exacerbations in symptomatology. Furthermore, the effect of antipsychotic medication on shape was also tested to determine if variable findings in previous work (Mamah et al., 2012; Torres et al., 2013) were due to differences in sex. Finally, the relationship between duration of illness and shape was tested to resolve whether deep brain regions are altered as a function of increased illness burden (Haijma et al., 2013).

## Materials and Methods

### Study Samples

This study included archival data on 6,861 participants from twenty-three cross-sectional study samples. In total, 2,826 individuals with schizophrenia (1,871 males and 955 females) and 4,035 healthy control participants (2,158 males and 1,877 females) were included in the meta-analysis, with data collected through the ENIGMA Schizophrenia Working Group. Measurement of clinical symptoms of the schizophrenia group was based on the Scale for the Assessment of Positive Symptoms (SAPS), the Scale for the Assessment of Negative Symptoms (SANS), and the Positive and Negative Symptom Scale (PANSS), as well as report of duration of illness and symptom severity (Andreasen, 1984; Andreasen, 1984; Kay & Opler, 1987). Participants provided written informed consent approved by local Institutional Review Boards prior to participating in their respective studies. The characteristics of this sample, including age, age of illness onset, duration of illness, and symptom severity, are provided in Table 1. A breakdown of the individual cohorts, including number of schizophrenia patients and control participants as well as age and sex, is provided in Table 2. The average age of schizophrenia participants was approximately 35.5 years (SD = 12.5), whereas the average age of healthy control participants was 33.5 years (SD = 13.0). The total scores across data sets were 63.9 for the PANSS (SD = 19.4), 15.9 for positive symptoms on the PANSS (SD = 5.8), 16.7 for negative symptoms on the PANSS (SD = 7.0), 16.5 for the SAPS (SD = 13.5), and 19.6 for the SANS (SD = 16.6). For data sets that recorded current antipsychotic type and dose, the average chlorpromazine (CPZ) dose was equivalent to 388.1 milligrams (Woods, 2003).

**Table 1.**
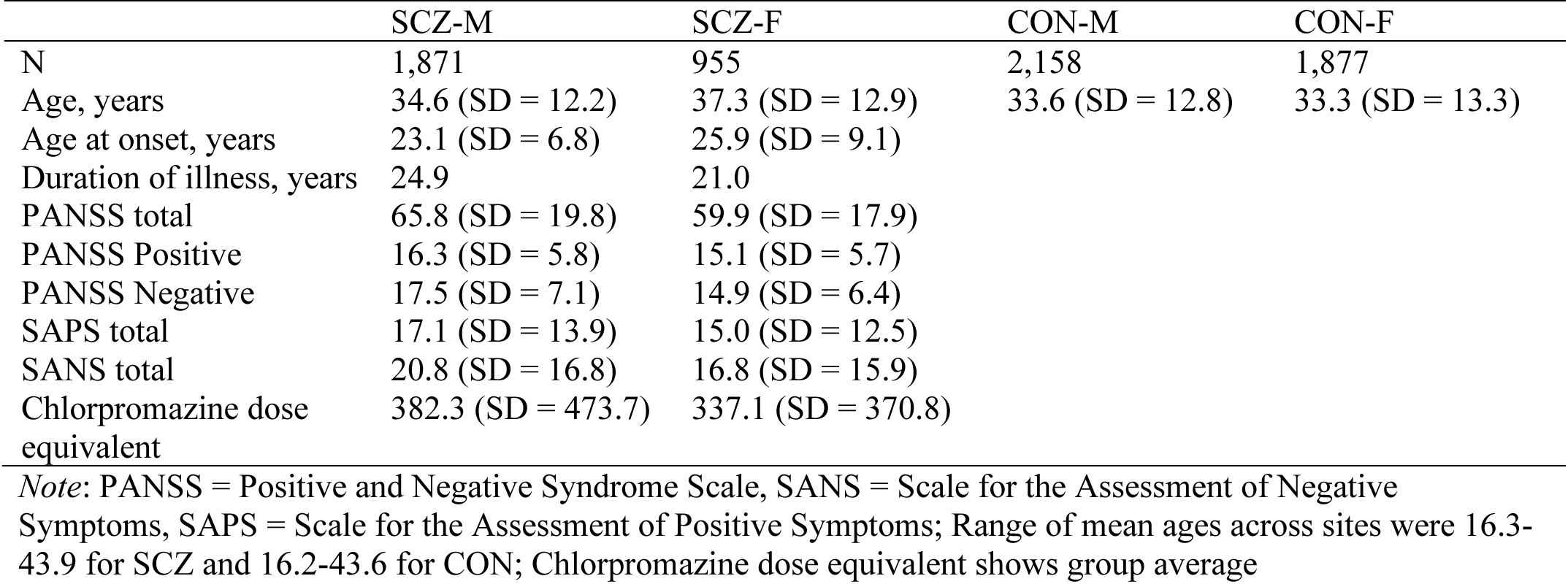
Sample characteristics.

**Table 2.** Sample Demographics by Cohort.

| Site | Schizophrenia |  |  | Healthy Control |  |  |
| --- | --- | --- | --- | --- | --- | --- |
|  | N | M/F | Age | N | M/F | Age |
| Basel SCORE<sup>1</sup> | 161 | 117/44 | 25.5 | 44 | 17/27 | 25.5 |
| CAMH<sup>2</sup> | 117 | 70/47 | 43.9 | 146 | 77/69 | 43.6 |
| Dublin<sup>1</sup> | 55 | 39/16 | 42.6 | 221 | 94/127 | 29.2 |
| FBIRN<sup>3</sup> | 185 | 139/46 | 38.9 | 174 | 124/50 | 37.5 |
| FIDMAG<sup>3</sup> | 160 | 124/36 | 39.6 | 123 | 54/69 | 37.5 |
| Galway<sup>1</sup> | 78 | 58/20 | 33.3 | 70 | 45/25 | 34.2 |
| Hubin<sup>1</sup> | 94 | 70/24 | 41.7 | 102 | 69/33 | 41.9 |
| Indiana | 227 | 180/47 | 37.8 | 496 | 352/144 | 35.7 |
| KASP SZ<sup>2</sup> | 56 | 34/22 | 30.3 | 32 | 15/17 | 27.5 |
| MPRC | 160 | 87/73 | 36.0 | 215 | 84/131 | 37.3 |
| Marburg<sup>1</sup> | 10 | 7/3 | 42.1 | 157 | 58/99 | 31.6 |
| NARSAD<sup>2</sup> | 26 | 17/9 | 26.6 | 69 | 45/24 | 27.3 |
| NU<sup>1</sup> | 108 | 74/34 | 34.1 | 92 | 51/41 | 31.9 |
| Olin | 302 | 168/134 | 37.8 | 1018 | 475/543 | 29.8 |
| Oxford<sup>2</sup> | 48 | 28/20 | 16.3 | 38 | 18/20 | 16.2 |
| PAFIP GE<sup>1</sup> | 142 | 88/54 | 29.7 | 79 | 49/30 | 27.7 |
| PAFIP Phillips<sup>1</sup> | 113 | 63/50 | 29.8 | 103 | 63/40 | 30.0 |
| StaLucia | 173 | 118/55 | 39.2 | 116 | 73/43 | 37.5 |
| TOP<sup>2</sup> | 219 | 130/89 | 32.0 | 303 | 159/144 | 35.4 |
| UCISZ<sup>3</sup> | 27 | 22/5 | 42.9 | 31 | 24/7 | 41.5 |
| UNINA<sup>2</sup> | 49 | 34/15 | 37.5 | 55 | 29/26 | 42.4 |
| UPenn<sup>1</sup> | 177 | 105/72 | 38.9 | 193 | 90/103 | 36.4 |
| Wellcome<sup>2</sup> | 61 | 41/20 | 27.7 | 88 | 48/40 | 30.2 |

<sup>1</sup> Sites used the SAPS/SANS to measure clinical symptoms (n = 1,310)
<sup>2</sup> Sites used the PANSS to measure clinical symptoms (n = 948)
<sup>3</sup> Sites used the SAPS/SANS and PANSS to measure clinical symptoms

### Image Acquisition and Processing

1. FreeSurfer Segmentation: High-resolution structural T1-weighted brain scans were acquired from each participant. Deep brain surface measures were characterized using the validated ENIGMA-Shape pipeline (Gutman et al., 2021). Deep brain regions of interest (ROI) were initially defined using the FreeSurfer segmentation protocol, which includes the thalamus, hippocampus, amygdala, pallidum, caudate, accumbens, and putamen from both the right and left hemispheres.
2. Surface Triangulation: In each subject, segmented ROIs were tessellated with a triangulated surface model with vertices that corresponded to a standard template surface (Wang et al., 2011). The resulting surfaces underwent visual quality control by individual raters at each participating site according to the ENIGMA-Shape Quality Control Guide and local quality control by the University of Southern California Imaging Genetics Center.
3. Surface Registration: The ENIGMA shape atlas provided a single template for all studies that contributed to the data set, averaging surface models of 200 unrelated individuals from the Queensland Twin Imaging Study.
4. Shape and Asymmetry Computation: Global brain volume measures (cortical gray matter, cerebral white matter, and cerebellar gray and white matter for covariate purposes), as well as deep brain shape deformation, were extracted from each individual. A medial curve was computed for individual shape models, and two measures of shape were calculated at each vertex: thickness was calculated as the distance to the medial curve, and the Jacobian determinant, known as “contraction” or “expansion,” was the ratio of the triangular area relative to that of the template at the corresponding vertices (Gutman et al., 2021; Wang et al., 2011). Group differences were assessed by comparing vertex-wise maps. To assess group differences in asymmetry between left and right hemispheres, asymmetry indices were computed and defined by the absolute difference between left and right thickness and surface expansion at each vertex.
5. Statistical Analyses: All statistical analyses were performed using the R statistical package (v4.2.2; R Core Team 2022). Vertex-wise mass univariate analysis per shape measure were performed first, followed by the aggregation of effect sizes, regression parameters, and confidence intervals for mass univariate meta-analysis. The meta-analysis was conducted by using the DerSimonian and Laird inverse-variance random-effects model as implemented in the R *metafor* package (Viechtbauer 2010). Linear model analyses were performed for participants with schizophrenia (SCZ) and healthy control participants (CON), comparing males (M) versus females (F) with and without a diagnosis of schizophrenia to assess for sex differences, as well as a broader comparison of schizophrenia patients versus healthy control participants per sex to determine the impact of psychosis on shape deformation of deep brain regions. To summarize, the following contrasts were conducted: 1) M/F; 2) DIAGNOSIS-by-SEX interaction; 3) SCZ-M/CON-M; 4) SCZ-F/CON-F; 5) SCZ-M/SCZ-F; and 6) CON-M/CON-F. Results from a SCZ/CON contrast were previously reported in a meta-analysis by our group (Gutman et al., 2022) that also used data from the ENIGMA Schizophrenia Working Group. Furthermore, twelve additional meta-regression models were run in SCZ-M and -F groups each to examine the relationships between shape deformation and clinical symptoms, medication use, or duration of illness (DURILL): 1) CPZ-M; 2) CPZ-F; 3) SAPS-M; 4) SAPS-F; 5) SANS-M; 6) SANS-F, 7) PANSSPOS-M; 8) PANSSPOS-F; 9) PANSSNEG-M; 10) PANSSNEG-F; 11) DURILL-M; and 12) DURILL-F. In all models, age was accounted for by including both its linear and quadratic terms, and intracranial volume (ICV) was also included as a covariate. Maps of p-values were corrected for multiple comparisons across all structures and measures for each linear model using a modified searchlight false discovery rate (FDR) procedure (Langers et al., 2007) then projected on a common reference surface.

## Results

### Main Effect of Diagnosis

Gutman and colleagues (2022) previously reported meta-analytic findings using data from the ENIGMA-Schizophrenia Working Group. Their results revealed more-concave-than-convex shape differences in the hippocampus, amygdala, nucleus accumbens, and thalamus in schizophrenia compared to healthy control participants. Furthermore, they highlighted more-convex-than-concave differences in the putamen and pallidum, with mixed findings in the caudate.

### Main Effect of Biological Sex

Sex differences in deep brain shape were observed in participants irrespective of diagnosis status (see Figure 1). Female brains exhibited a pattern of lower thickness and greater surface contraction in the bilateral thalamus, pallidum, putamen, and amygdala compared to males. Findings were mixed in the hippocampus and caudate. There were subtle patterns of surface contraction in the accumbens. The magnitude of the above effects, including thickening/thinning (*d* = -0.30 to 0.20) and surface expansion/contraction (*d* = -0.35 to 0.15), ranged from small to moderate. See Table 3 for percentage of surface area affected.

**Figure 1.**
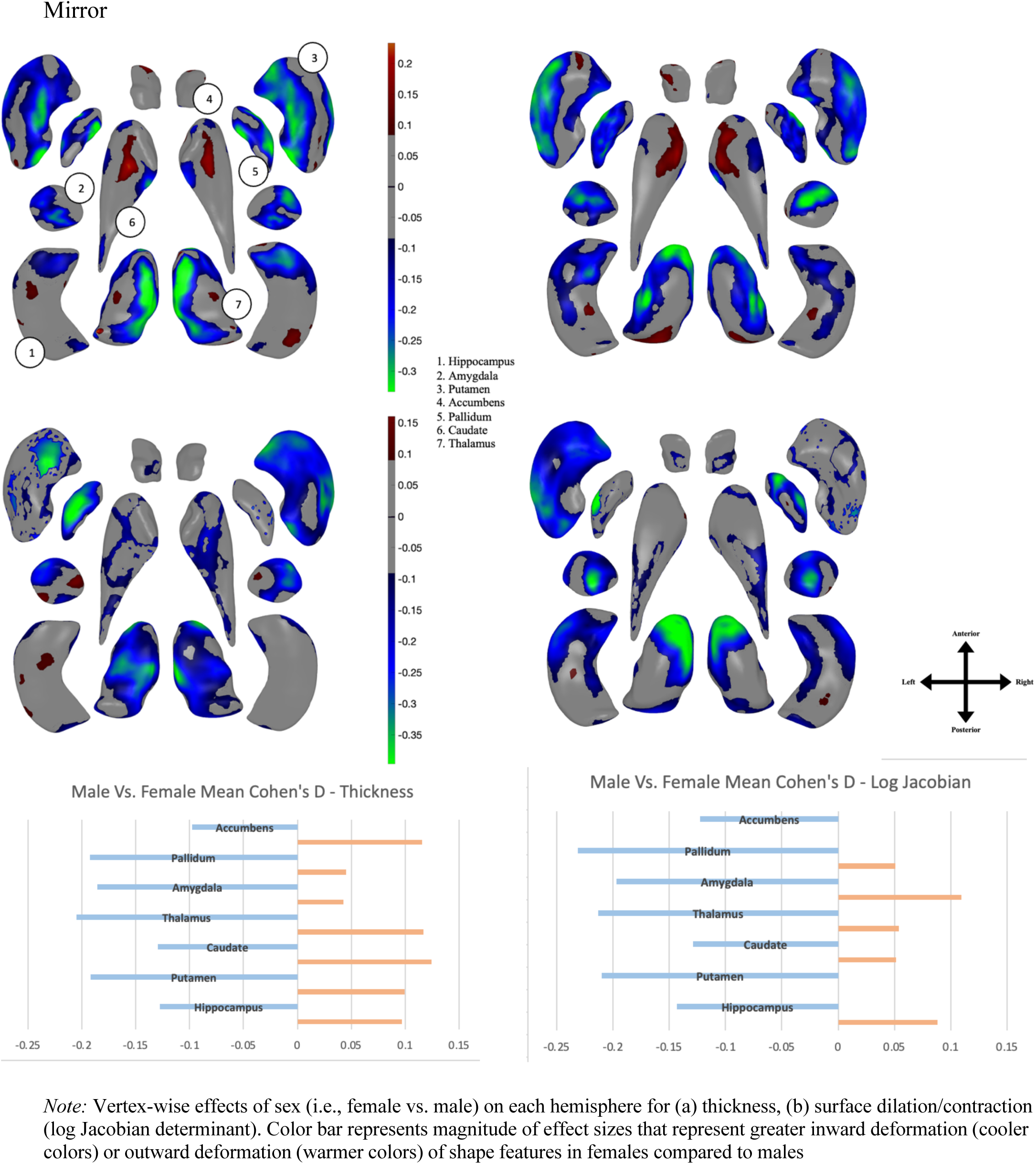
Deep Brain Shape Deformation in Females vs. Males.

**Table 3.**
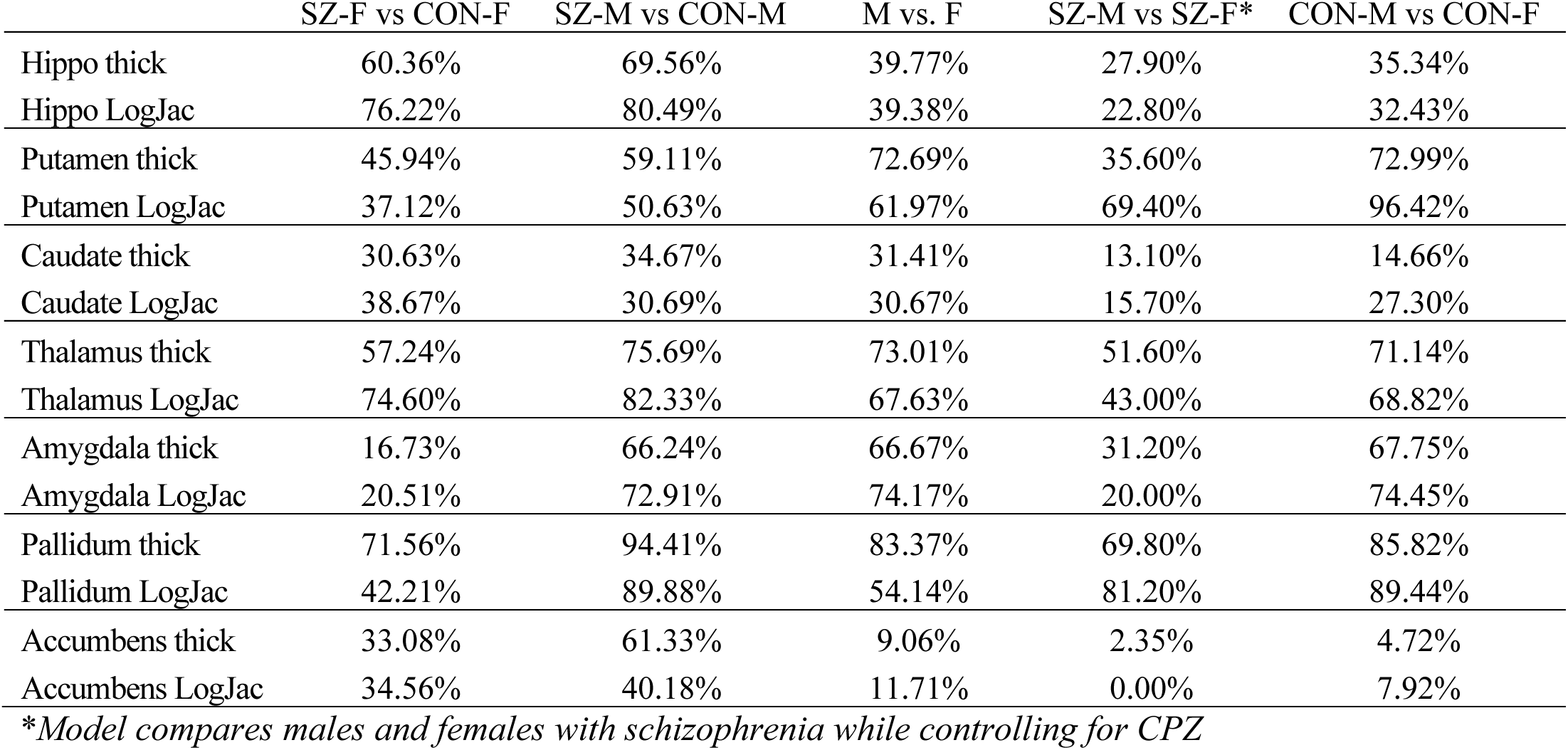
Percentage Surface Area Affected Across Models.

The Diagnosis-by-Sex interaction models were non-significant for all tested regions suggesting a diagnosis of schizophrenia does not meaningfully contribute to differential patterns of shape deformation between males and females.

### Effect of Diagnosis on Bilateral Shape Features in Males

Patterns of deep brain shape deformation were observed across several regions based on diagnosis in male participants (see Figure 2). Within the bilateral hippocampus, amygdala, accumbens, and thalamus, there was a pattern of significant thinning and surface contraction in males with schizophrenia compared with control participants. Within the bilateral putamen and pallidum, there was a pattern of significant thickening and surface expansion in males with schizophrenia. Findings were mixed for the caudate, with increased thinning observed in the medial caudate and more thickening in the lateral regions of the caudate. The magnitude of the above effects, including thickening/thinning (*d* = -0.30 to 0.30) and surface expansion/contraction (*d* = -0.25 to 0.25), ranged from small to moderate. See Table 3 for percentage of surface area affected.

**Figure 2.**
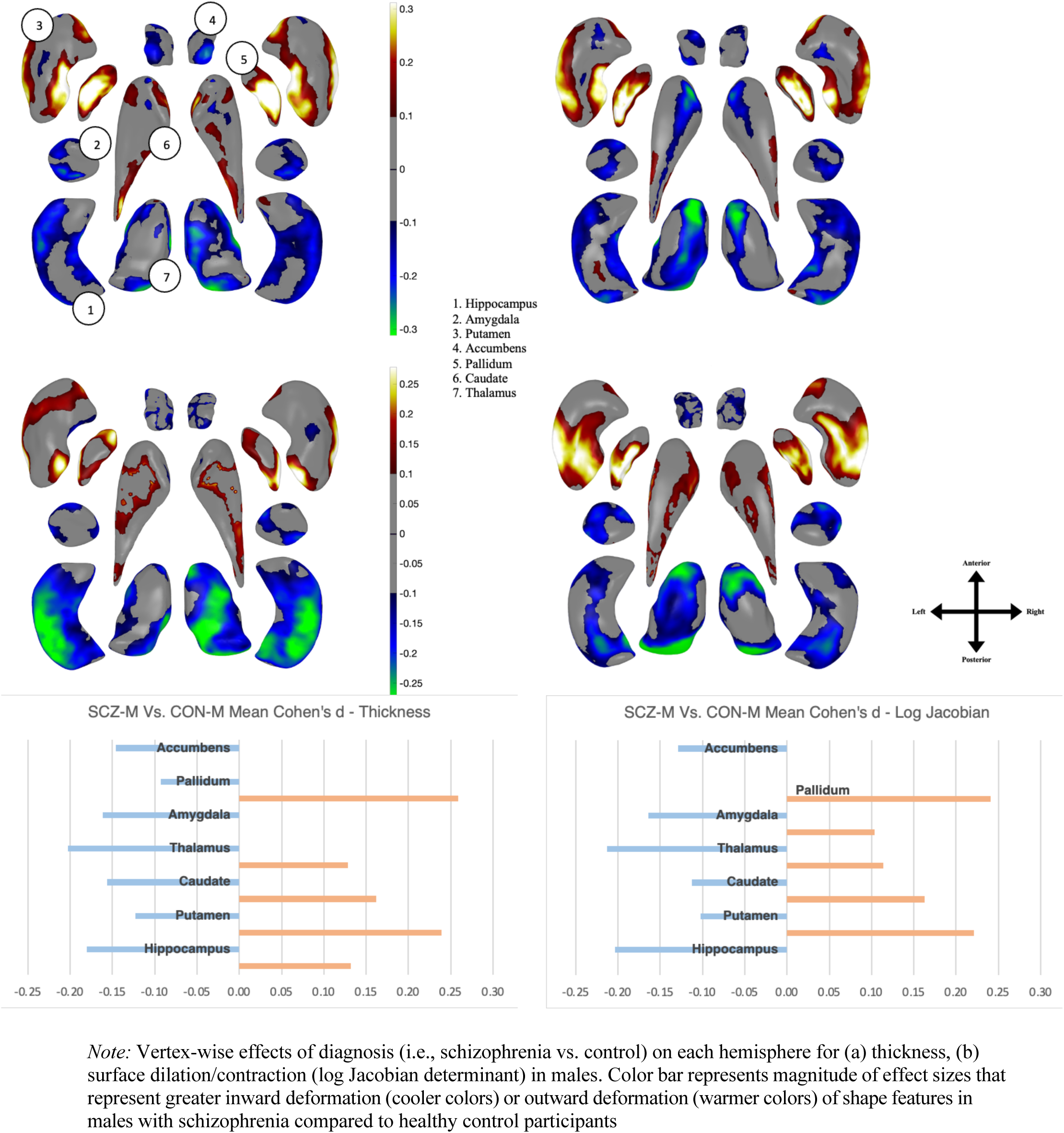
Deep Brain Shape Deformation in SCZ Males vs. CON Males.

### Effect of Diagnosis on Bilateral Shape Features in Females

Patterns of deep brain shape deformation were also observed across multiple structures based on diagnosis in female participants (see Figure 3). Within the bilateral hippocampus, amygdala, accumbens, and thalamus, there was a pattern of significant thinning and surface contraction in females with schizophrenia compared with control participants. Within the bilateral putamen and pallidum, there was a pattern of significant thickening and surface expansion in females with schizophrenia. Similar to males, findings were mixed when it came to the caudate, with greater thinning in the medial caudate and more thickening in the lateral regions. The magnitude of the above effects, including thickening/thinning (*d* = -0.30 to 0.30) and surface expansion/contraction (*d* = -0.30 to 0.30), ranged from small to moderate. See Table 3 for percentage of surface area affected.

**Figure 3.**
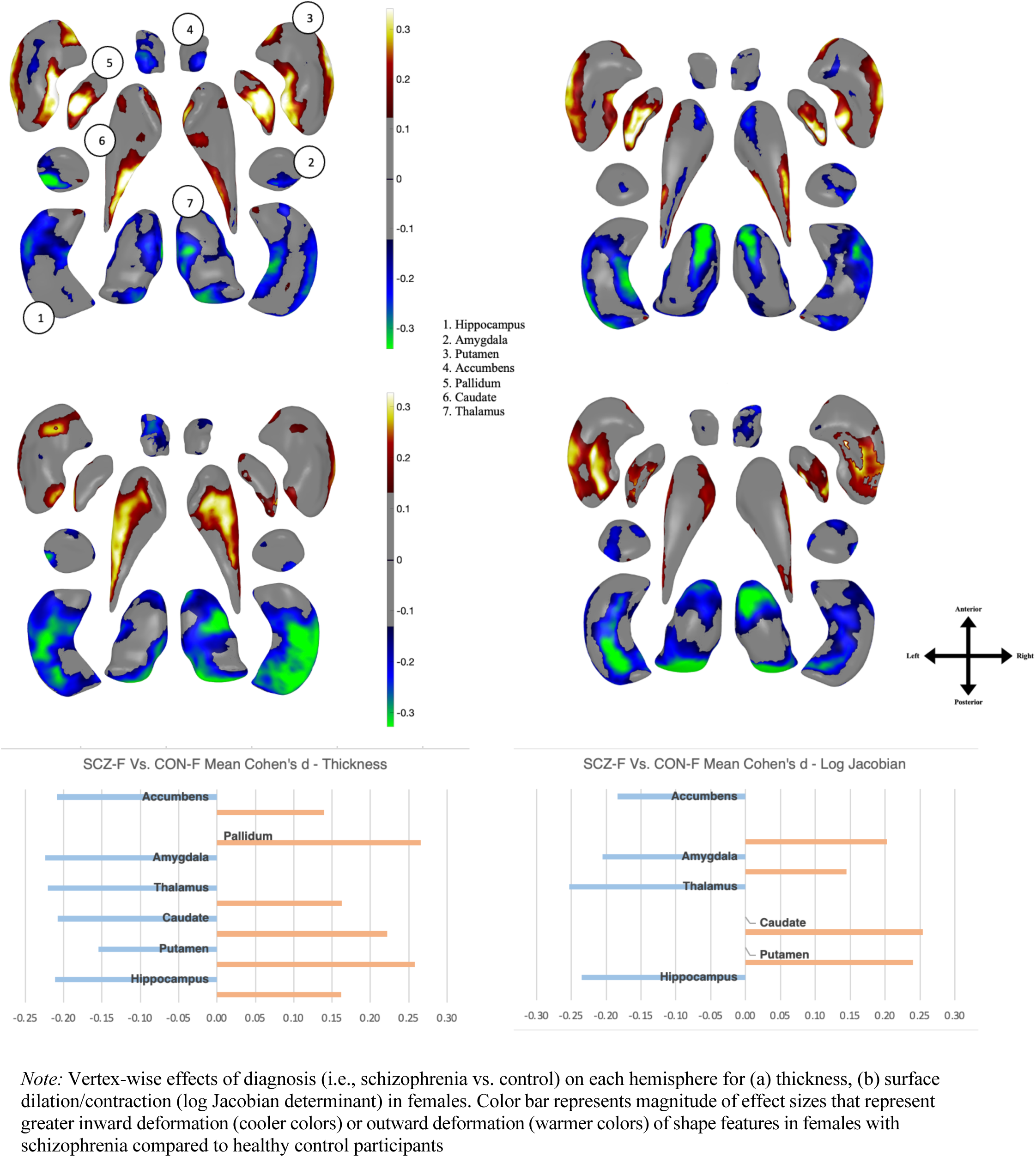
Deep Brain Shape Deformation in SCZ Females vs. CON Females.

### Effect of Sex on Bilateral Shape Features in Healthy Control Participants

Patterns of deep brain shape deformation were observed across all structures on the basis of biological sex in healthy control participants (see Figure 4). Within the bilateral thalamus, pallidum, putamen, and amygdala, there was a pattern of significant thinning and surface contraction in females compared to male control participants. Findings were mixed in the bilateral hippocampus and caudate, with medial regions of the caudate showing greater thickness in females compared to males, and anterior regions of the hippocampus showing more thinning in females. Greater surface expansion was observed in medial regions of the hippocampus in females compared to males, with surface contraction in the tail of the caudate. Overall, the most extensive shape differences were observed in the dorsal putamen, amygdala, left pallidum, and dorsal thalamus, with none observed in the accumbens. The magnitude of the above effects, including thickening/thinning (*d* = -0.30 to 0.30) and surface expansion/contraction (*d* = -0.30 to 0.20) ranges from small to moderate.

**Figure 4.**
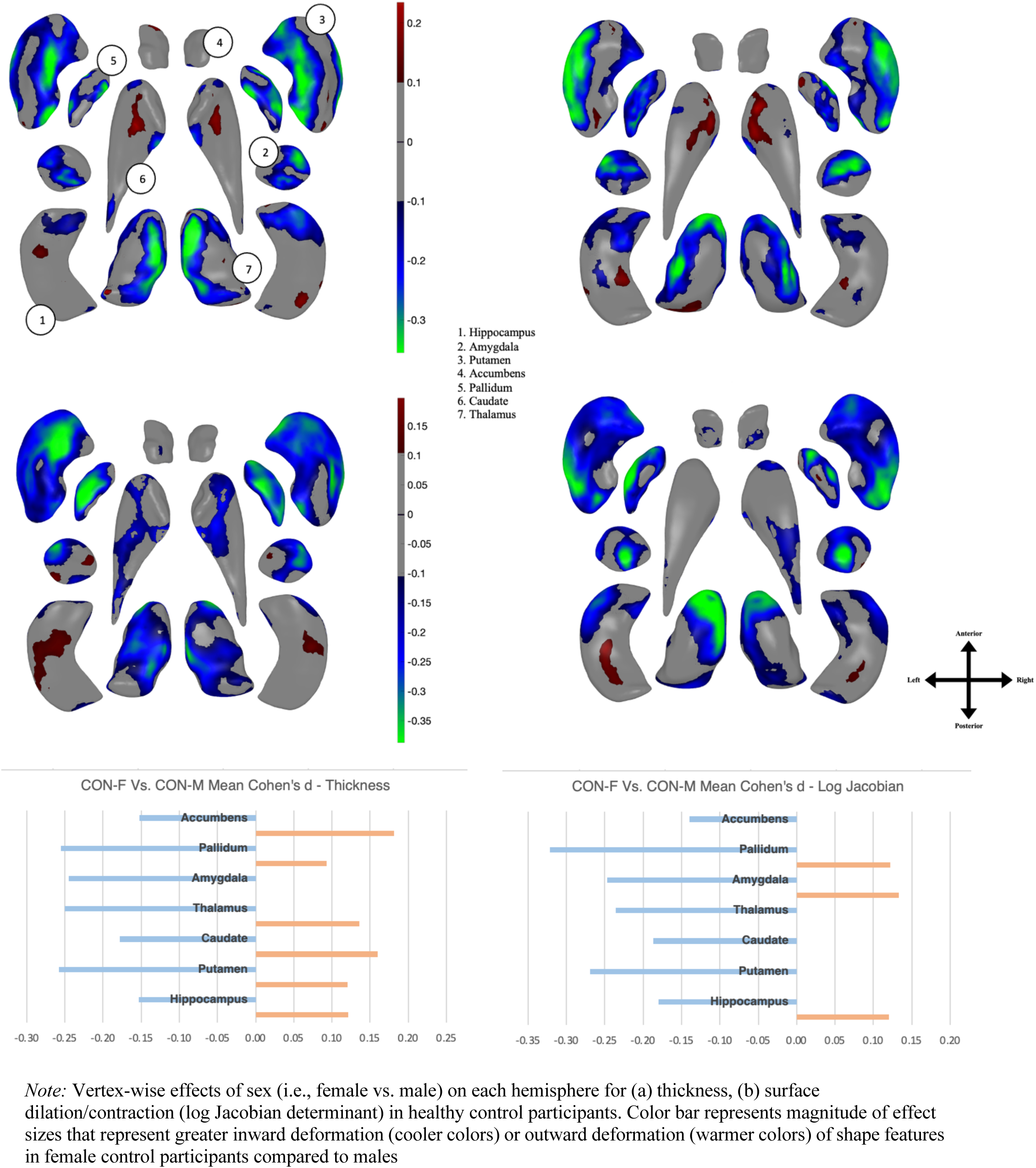
Deep Brain Shape Deformation in CON Females vs. CON Males.

### Effect of Sex on Bilateral Shape Features in Participants With Schizophrenia

Patterns of deep brain shape deformation were also observed across all structures among males and females with schizophrenia (see Figure 5). Of note, this model controlled for chlorpromazine (CPZ) dose equivalents. Within the bilateral pallidum, putamen, thalamus, and amygdala, there was a pattern of significant thinning and surface contraction in females compared to males with schizophrenia. Similar to healthy control participants, anterior regions of the hippocampus showed more thinning in females. Medial regions of the caudate showed greater thickness in females compared to males. Overall, the most extensive patterns of deformation were observed in dorsal, ventral, and lateral aspects of putamen, thalamus, amygdala, and pallidum. The magnitude of the effects, including thickening/thinning (*d* = -0.30 to 0.30) and surface expansion/contraction (*d* = -0.30 to 0.20), ranged from small to moderate.

**Figure 5.**
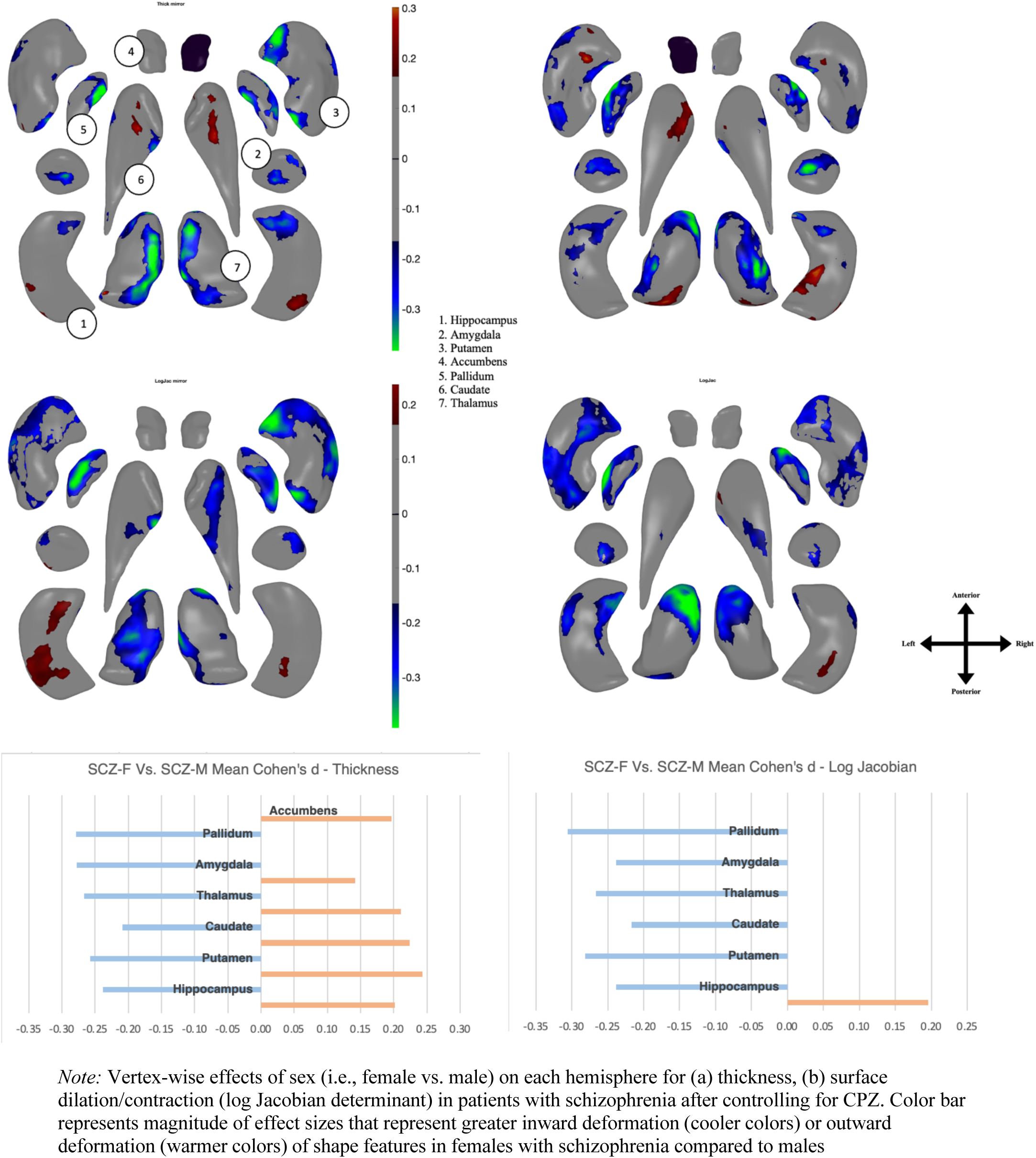
Deep Brain Shape Deformation in SCZ Females vs. SCZ Males.

### Effect of Diagnosis on Shape Asymmetry in Males

Across all regions of interest, the asymmetry index was defined as the absolute difference in effect size between both hemispheres. Of note, only 21 sites provided unilateral data for left and right hemispheres and, therefore, two sites were excluded from asymmetry analyses. Small, but significant, group differences in asymmetry patterns were observed in males with schizophrenia compared to healthy control participants (see Figure 6). Regarding thickness, the asymmetry index was greater for men with schizophrenia as compared to healthy control participants for the hippocampus and accumbens (*d* = 0.05 to 0.15). The asymmetry index was smaller for the thalamus (*d* = -0.05 to -0.20). Group differences in the asymmetry index were negligible for the amygdala, caudate, putamen, and pallidum. Regarding surface expansion, the asymmetry index was greater for men with schizophrenia as compared to healthy control participants for the hippocampus, thalamus, and accumbens (*d* = 0.05 to 0.15). For the amygdala, pallidum, caudate, and putamen, group differences were negligible. The magnitude of the above effects is small.

**Figure 6.**
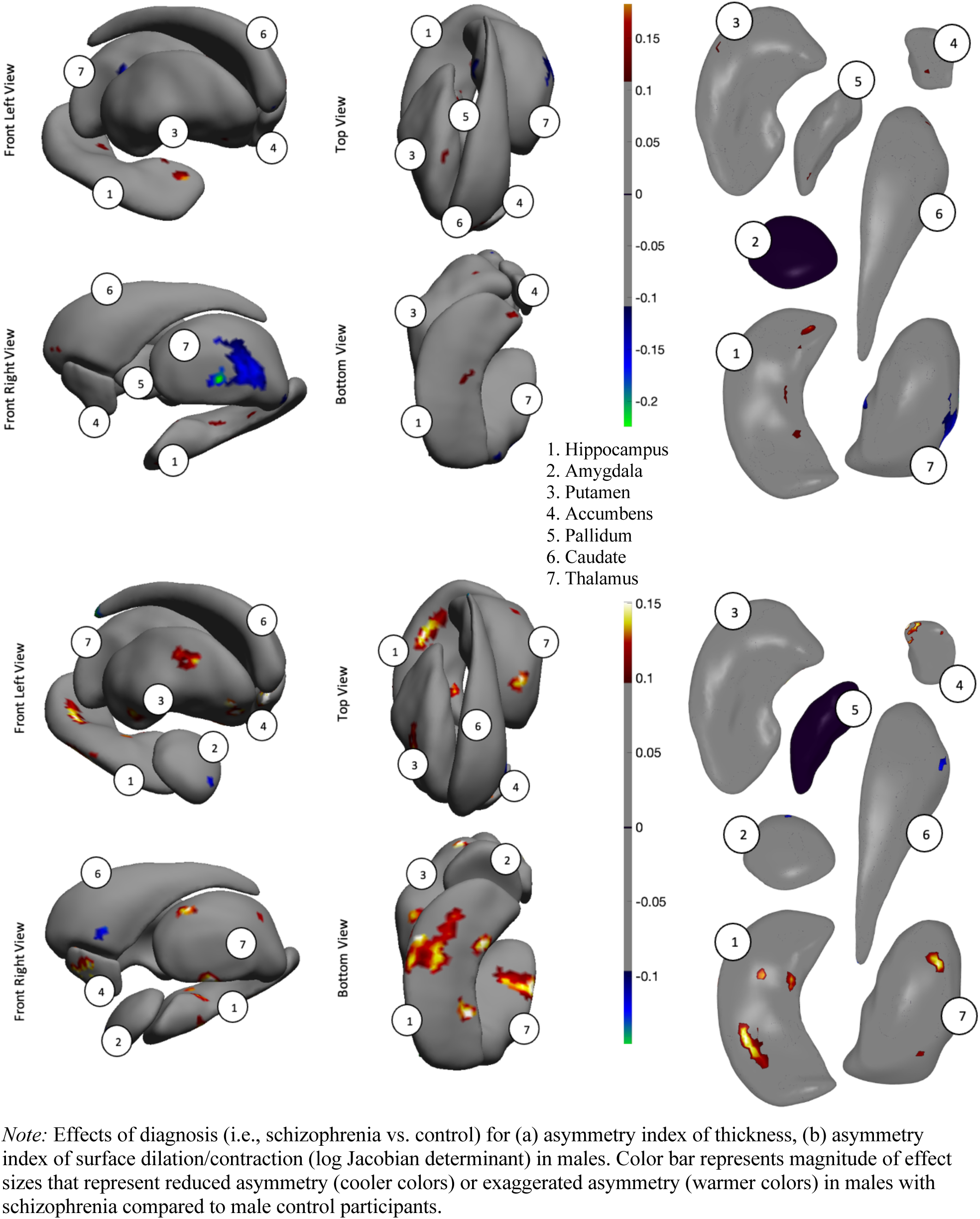
Effect of Diagnosis on Asymmetry Index for SCZ Males vs. CON Males.

### Effect of Diagnosis on Shape Asymmetry in Females

Small, but significant, group differences in asymmetry patterns were also observed in females with schizophrenia compared to healthy control participants (see Figure 7). Regarding thickness, the asymmetry index was smaller for women with schizophrenia as compared to healthy control participants for the thalamus (*d* = -0.05 to -0.15). Group differences in the asymmetry index were negligible for all other regions of interest. Regarding surface expansion, the asymmetry index was greater for women with schizophrenia as compared to healthy control participants for the putamen and thalamus (*d* = 0.05 to 0.15). For the hippocampus, amygdala, pallidum, caudate, and accumbens, group differences were negligible. The magnitude of the above effects is small.

**Figure 7.**
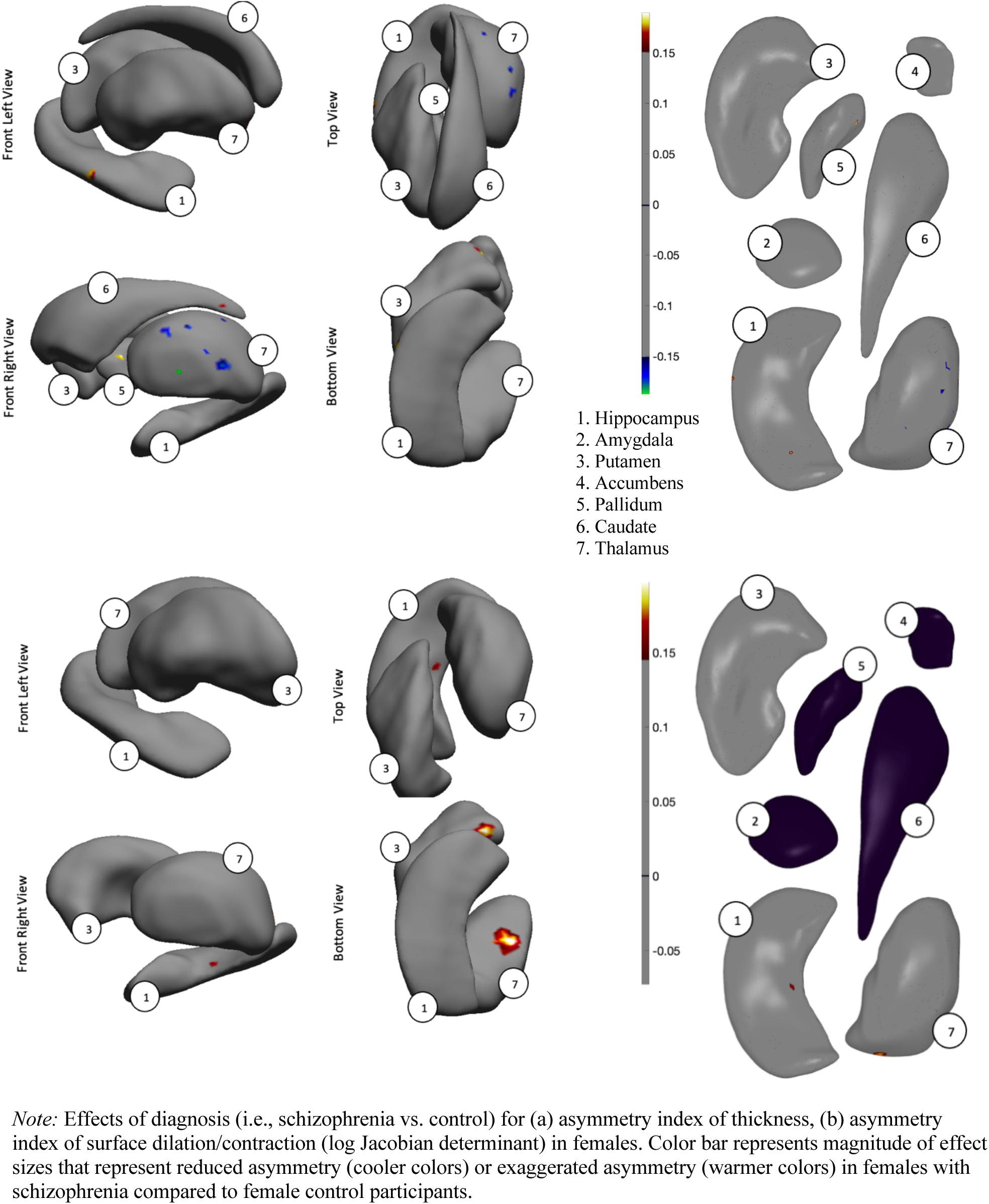
Effect of Diagnosis on Asymmetry Index for SCZ Females vs. CON Females.

### Effect of Sex on Shape Asymmetry in Healthy Control Participants

In healthy control participants, small but significant group differences in asymmetry patterns were observed in females compared to males (see Figure 8). Regarding thickness, the asymmetry index was smaller for female compared to male control participants for the amygdala, pallidum, and thalamus (*d* = -0.05 to -0.15). Group differences in the asymmetry index were negligible for the hippocampus, caudate, accumbens, and putamen. Regarding surface expansion, the asymmetry index was greater for female compared to male control participants for the thalamus, amygdala, and caudate (*d* = -0.05 to -0.15). For the hippocampus, pallidum, accumbens, and putamen, group differences were negligible. The magnitude of the above effects is small.

**Figure 8.**
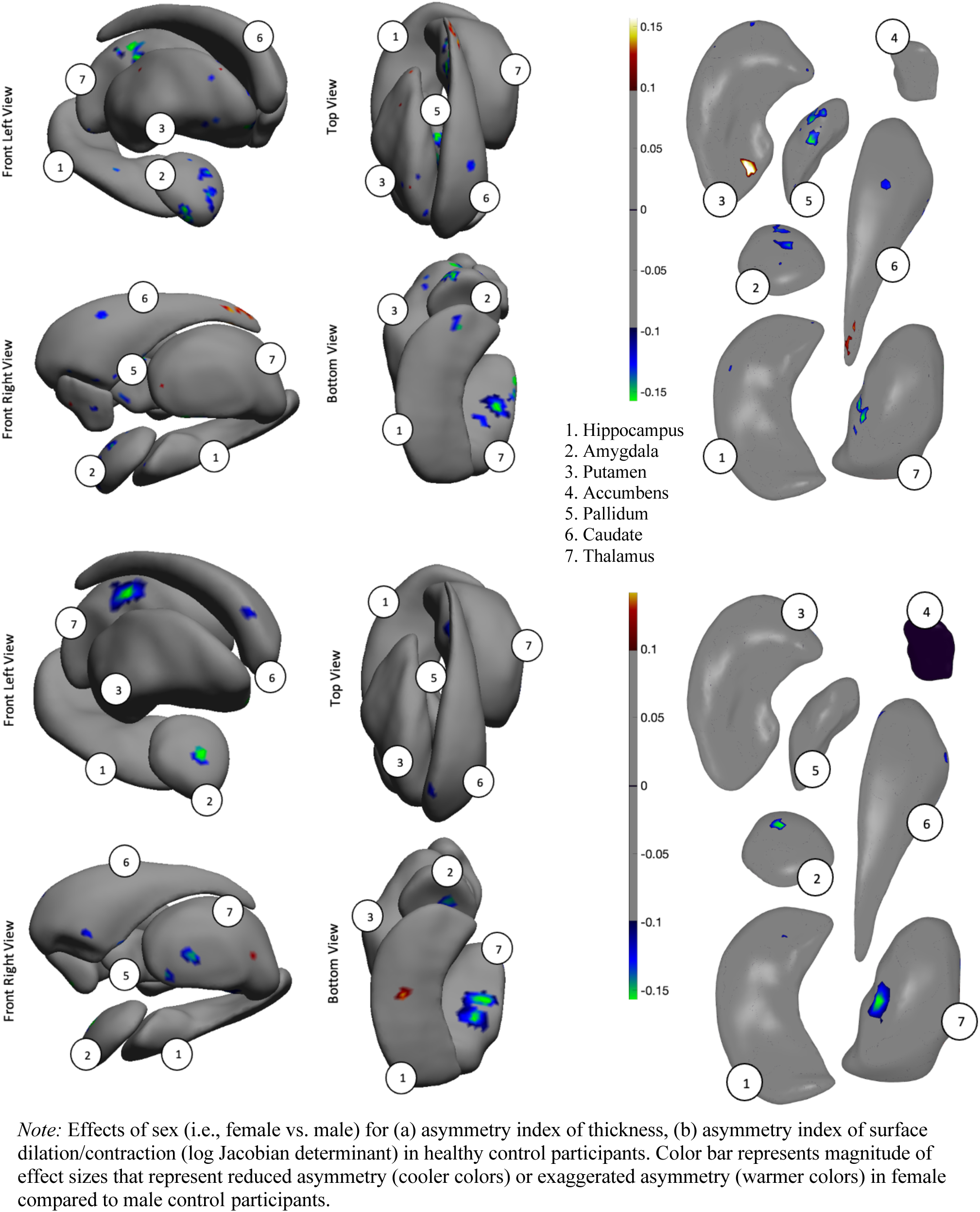
Effect of Sex on Asymmetry Index for CON Females vs. CON Males.

### Effect of Sex on Shape Asymmetry in Schizophrenia

In patients with schizophrenia, few group differences in asymmetry patterns were observed in females compared to males (see Figure 9). Regarding thickness, the asymmetry index was smaller for female compared to male patients for the caudate and putamen (*d* = -0.05 to -0.12). Group differences in the asymmetry index were negligible for the remaining regions of interest. Regarding surface expansion, group differences in the asymmetry index were negligible across all regions of interest. The magnitude of the above effects is small.

**Figure 9.**
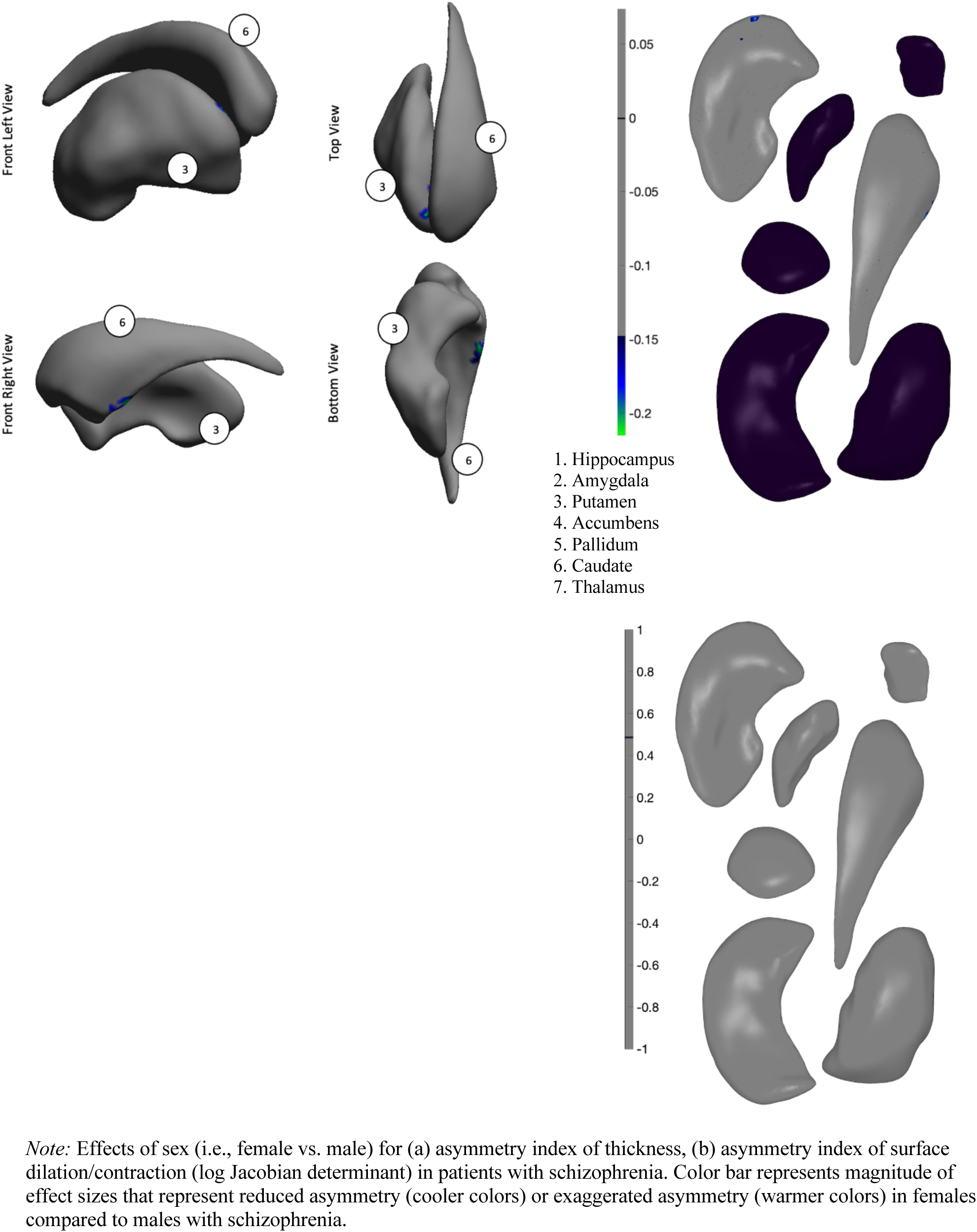
Effect of Sex on Asymmetry Index for SCZ Females vs. SCZ Males.

### Clinical Correlates in Male Schizophrenia Patients

Meta-regression models were run to examine correlations between shape deformation and clinical symptoms, as well as medication use. Only 13 sites provided chlorpromazine dose equivalents (CPZ), 12 sites provided data on participants’ SAPS/SANS scores, 10 sites provided participants’ PANSS scores, and 17 sites provided data on duration of illness. The remaining sites were not included in these regression models. Using shape meta-analysis, there were small but statistically significant relationships between higher CPZ and shape-derived thinning (*d* = -0.0002 to 0.0004) and surface contraction (*d* = -0.00004 to 0.00004) in the caudate, accumbens, hippocampus, amygdala, and thalamus (see Figure 10). In addition, there were small but statistically significant relationships between elevated SAPS scores (i.e., positive symptoms) and shape-derived thinning (*d* = -0.006 to -0.003) and surface contraction (*d* = -.0016 to -.0007) in the bilateral caudate, right hippocampus, and right amygdala (see Figure 12). After FDR correction, there were no significant relationships between SANS scores (i.e., negative symptoms), PANSSPOS scores, PANSSNEG scores, or duration of illness with deep brain shape deformation.

**Figure 10.**
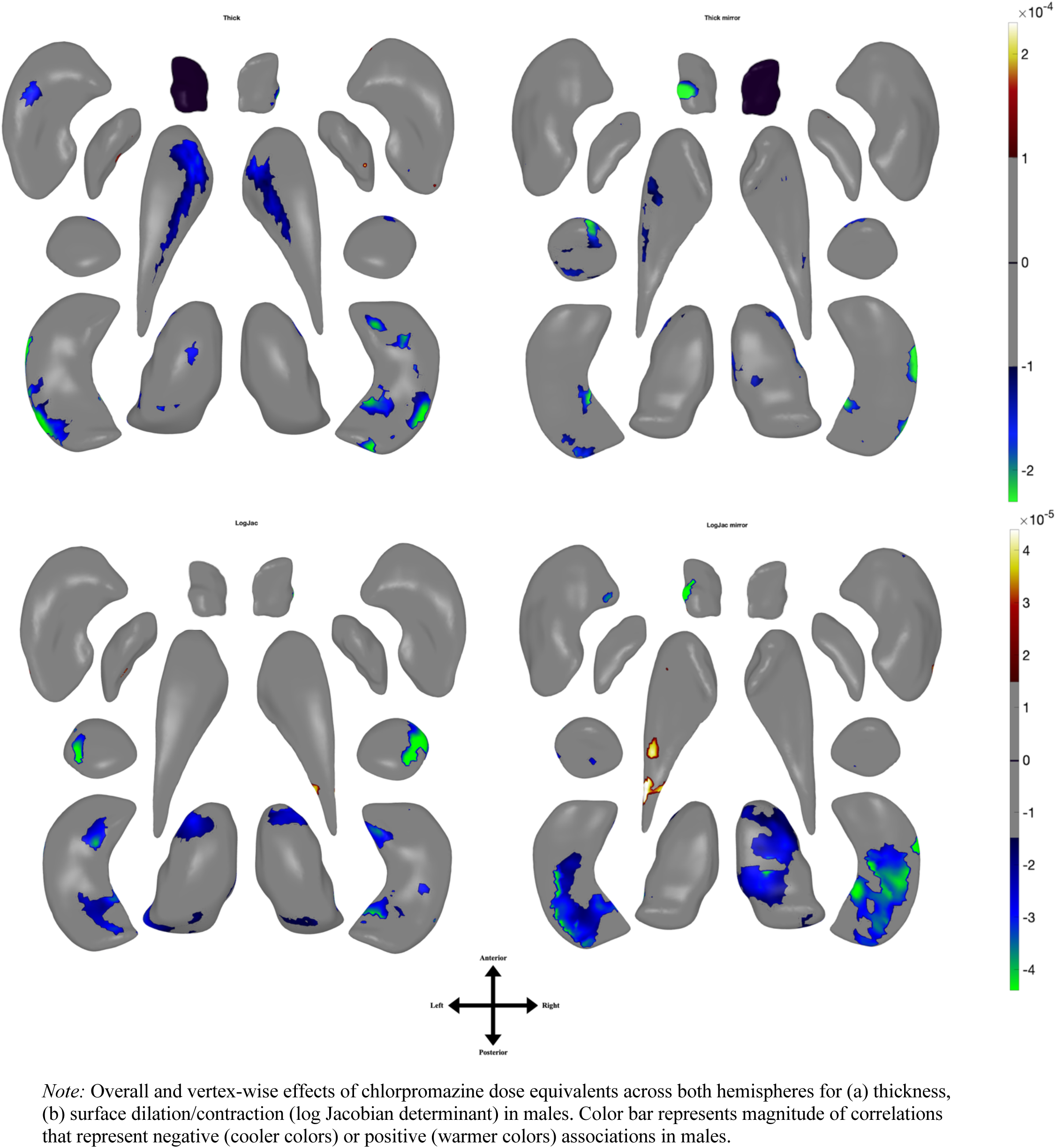
Effects of CPZ on Deep Brain Shape Deformation in Males.

### Clinical Correlates in Female Schizophrenia Patients

In female patients shape meta-analysis meta-regression revealed small but statistically significant relationships between increased CPZ and shape-derived thinning (*d* = -0.0002 to 0.0004) and surface contraction (*d* = -0.00006 to 0.00006) in caudate, accumbens, hippocampus, amygdala, and thalamus (see Figure 11). In addition, there were small but statistically significant relationships between elevated SAPS scores and shape-derived thinning (*d* = -0.006 to -0.003) and surface contraction (*d* = -0.0016 to -0.0007) in the bilateral caudate, right hippocampus, and right amygdala (see Figure 13). There were no significant relationships between SANS scores, PANSSPOS scores, PANSSNEG scores, or duration of illness and shape deformation.

**Figure 11.**
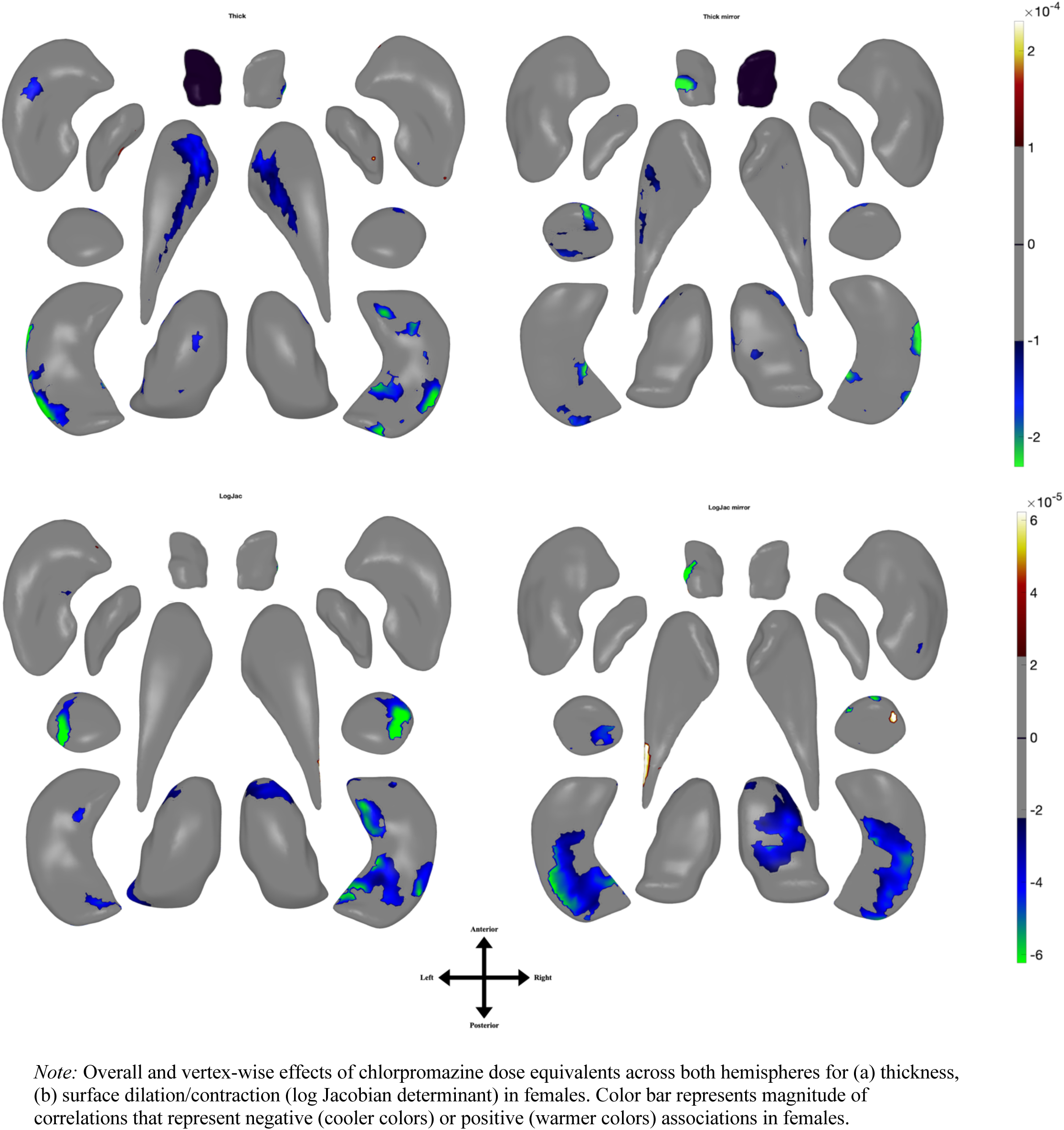
Effects of CPZ on Deep Brain Shape Deformation in Females.

**Figure 12.**
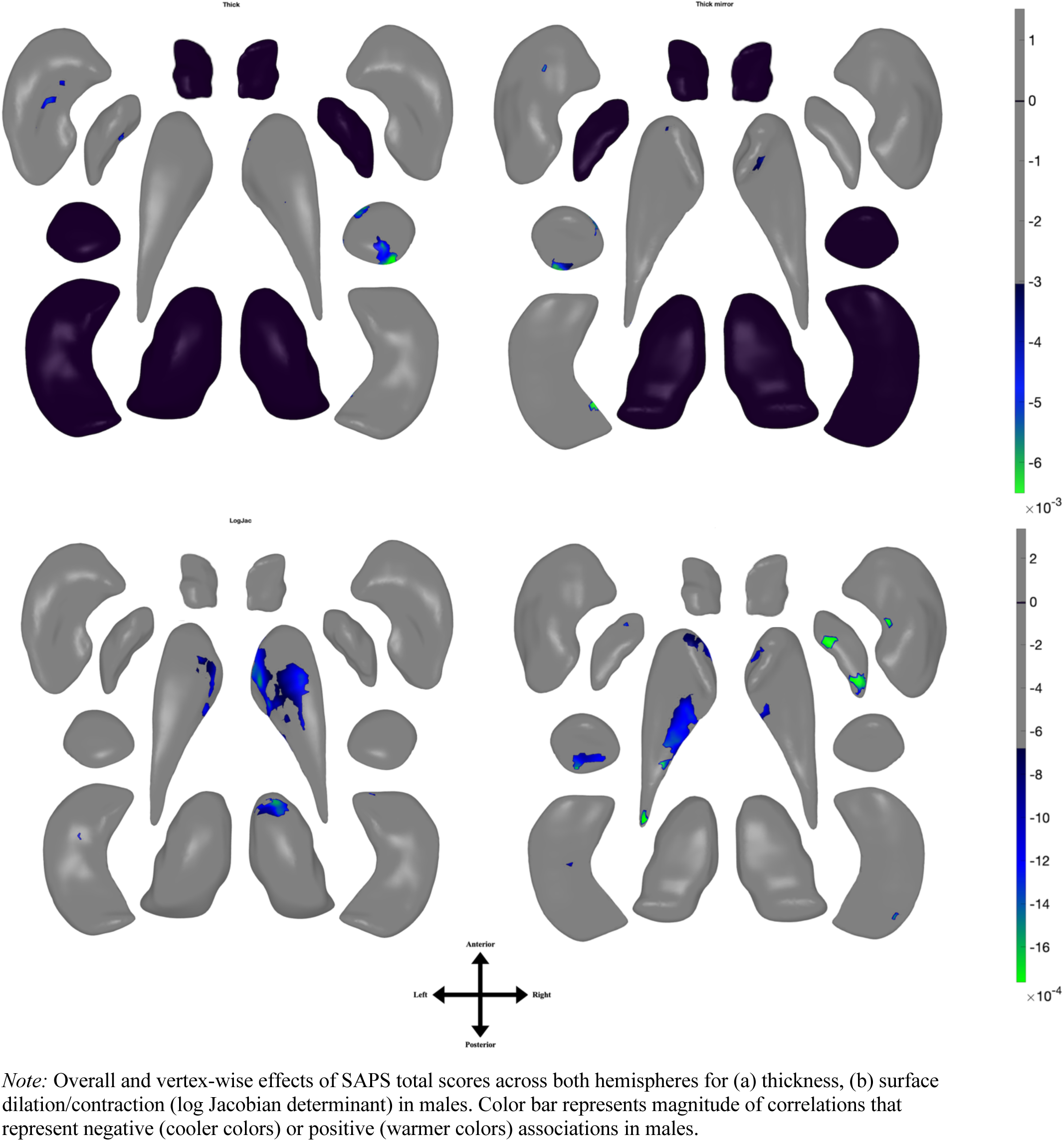
Effects of SAPS on Deep Brain Shape Deformation in Males.

**Figure 13.**
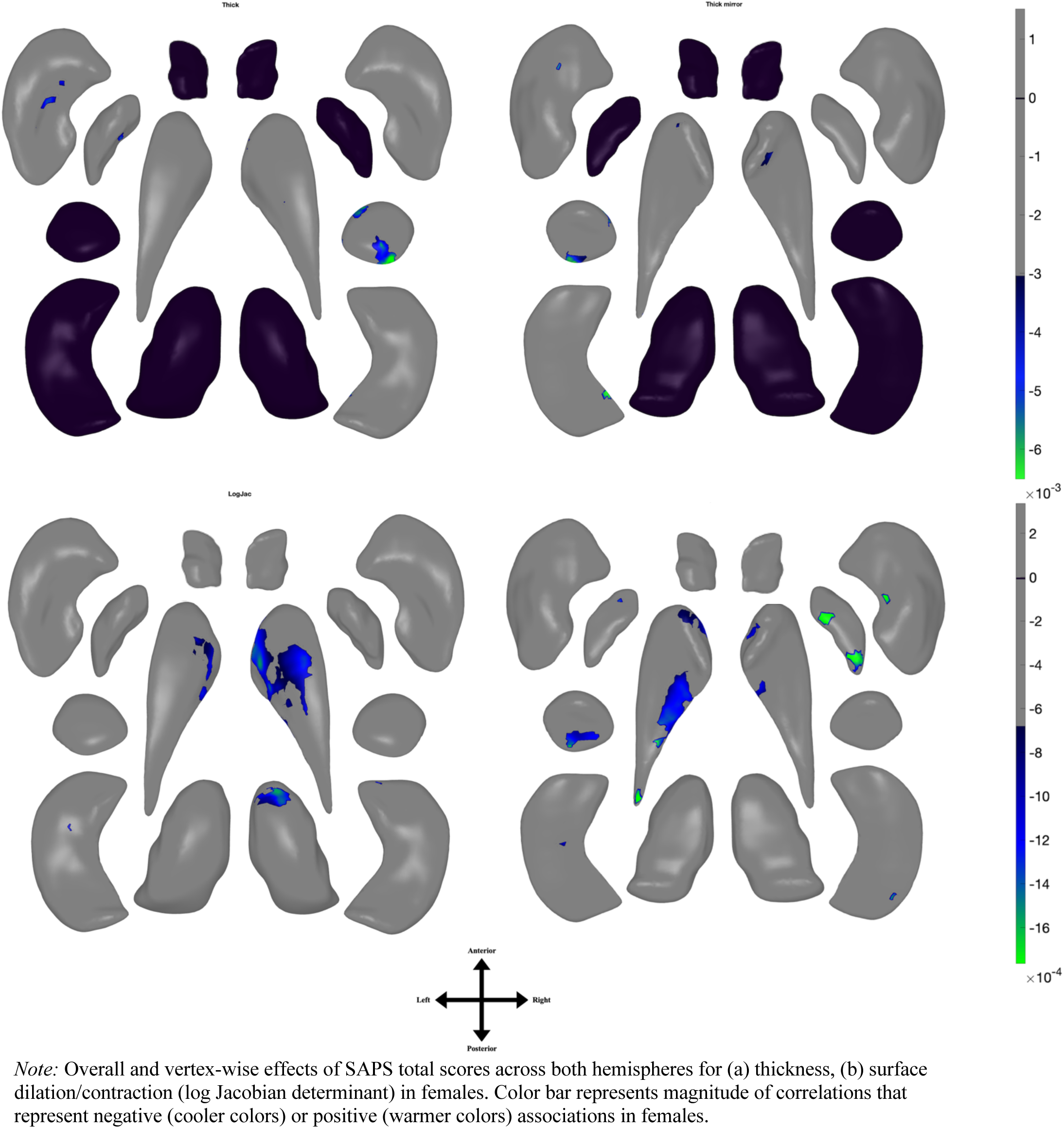
Effects of SAPS on Deep Brain Shape Deformation in Females.

Regarding male/female patient comparisons of clinical correlates, there was a slightly greater percent of affected surface area in the putamen, caudate, and thalamus in males compared to females for the CPZ models. For SAPS models, slightly greater affected surface area was observed only in the amygdala among male versus female patients (5.92% versus 5.83%); see Table 4 for all comparisons.

**Table 4.** Percentage Surface Area Affected Across Clinical Correlates.

|  | CPZ_Males | CPZ_Females | SAPS_Males | SAPS_Females |
| --- | --- | --- | --- | --- |
| Hippo thick | 10.96% | 10.54% | 0.16% | 0.16% |
| Hippo LogJac | 20.10% | 21.65% | 0.26% | 0.26% |
| Putamen thick | 0.73% | 0.69% | 0.57% | 0.57% |
| Putamen LogJac | 0.30% | 0.29% | 0.11% | 0.11% |
| Caudate thick | 12.61% | 12.31% | 0.77% | 0.77% |
| Caudate LogJac | 1.05% | 0.67% | 14.53% | 14.53% |
| Thalamus thick | 4.91% | 4.38% | 0.00% | 0.00% |
| Thalamus LogJac | 19.85% | 12.72% | 1.12% | 1.12% |
| Amygdala thick | 9.48% | 9.29% | 5.92% | 5.83% |
| Amygdala LogJac | 7.59% | 13.22% | 2.17% | 2.17% |
| Pallidum thick | 0.55% | 0.55% | 0.76% | 0.76% |
| Pallidum LogJac | 0.10% | 0.00% | 2.21% | 2.21% |
| Accumbens thick | 5.50% | 5.50% | 0.00% | 0.00% |
| Accumbens LogJac | 2.50% | 2.50% | 0.00% | 0.00% |

## Discussion

The overarching aim of this study was to conduct a meta-analysis of sex differences in deep brain shape deformation among patients with schizophrenia compared to healthy individuals. Group differences and asymmetry indices were examined between males vs. females (irrespective of diagnostic status); males with schizophrenia vs. healthy males; females with schizophrenia vs. healthy females; males vs. females with schizophrenia; and male vs. female healthy individuals. An overall model assessing the interaction between diagnosis and sex in patterns of shape deformation was also conducted. Finally, correlates of clinical symptomatology, illness duration, and medication use with deep brain shape deformation were also examined in schizophrenia patients.

Given the literature on brain pathways and functional impairment in schizophrenia, it was hypothesized that participants with schizophrenia would exhibit abnormal patterns of shape deformation in the amygdala, hippocampus, thalamus, and basal ganglia (i.e., caudate, putamen, globus pallidus, and nucleus accumbens) compared to healthy control participants. Furthermore, it was hypothesized that sex differences exist in the brain, such that males would demonstrate greater less concave shape in the hippocampus, amygdala, and putamen compared to females, with females exhibiting less concave shape in the thalamus compared to males. Findings from the present study provide partial support for these hypotheses, primarily revealing significant main effects for sex on deep brain shape deformation, or morphology. The diagnosis-by-sex interaction model revealed null findings for all regions, suggesting that schizophrenia does not have an appreciable differential impact on normal sex differences in deep brain nuclei shape. Despite some variability relative to previous work, this project is the first to explicitly study sex-based structural differences in schizophrenia from a meta-analytic perspective using high-dimensional brain mapping procedures, and thus leverages the strengths of large and diverse samples to powerfully inform questions at the intersection of sex and disease.

The findings of the present study are largely consistent with prior neuroimaging deep brain analyses of patients with schizophrenia compared to healthy individuals (Gutman et al., 2022; Okada et al., 2016; Tu et al., 2022). In both male and female patients with schizophrenia, there was a predominance of regions with more-concave-than-convex shape differences across the bilateral hippocampus, amygdala, accumbens, and thalamus, with more-convex-than-concave differences in the putamen and pallidum in both male and female comparisons of SCZ/CON. Our previous work on surface-based shape in deep brain regions (Gutman et al., 2022) revealed greater shape deformation in the bilateral hippocampus, amygdala, thalamus, and accumbens compared to healthy control participants, whereas surface expansion was reported in the bilateral putamen, pallidum, and caudate. These results match strongly with the male and female SCZ vs. CON models in the present study, which showed modest variation in the pattern, but consistent in region and direction of effect. As representations of localized volume loss, our shape deformation findings also find agreement with volume-based work. An additional study using estimates of subcortical volumes from ENIGMA schizophrenia sites (van Erp et al., 2016) revealed markedly similar SCZ vs. CON pattern with Gutman and colleagues (2022) for the hippocampus, amygdala, thalamus, and accumbens. In addition, they observed that sex differences might influence the direction and magnitude of these subcortical volumes given they found that differences in sample proportions of males for the sites had negative associations with the accumbens and amygdala effect sizes, suggesting a basis for differential sex-based patterns of brain abnormalities (van Erp et al., 2016). In a meta-analysis of 2,564 participants, decreased volume in the bilateral hippocampus, amygdala, thalamus, and accumbens was observed in patients with schizophrenia compared to healthy control participants, whereas increased volumes were observed in the bilateral caudate, putamen, and pallidum (Okada et al., 2016). Furthermore, a single-site study of 160 participants with schizophrenia and 160 healthy control participants reported a significant reduction in hippocampal and thalamic volumes, along with increased volume of the pallidum (Tu et al., 2022).

The sex-based pattern and direction of surface-based deformation observed when testing the main effects of sex was one of the most consistent findings across all of the sex-focused contrasts. This consisted of reduced thickness/surface-contraction (i.e., concave deformation) in the amygdala, thalamus, putamen, pallidum and accumbens in women, and mixed findings (both concave and convex deformation) in the caudate and hippocampus. The same pattern emerged in healthy females compared to healthy males, with the greatest reductions in thickness/surface expansion in the dorsal putamen, amygdala, left pallidum, and dorsal thalamus, and mixed directions in the hippocampus and caudate. Shape deformation between men and women with schizophrenia were also markedly similar to the main effect of sex contrast, albeit with subtle shifts in the pattern, highlighting the durability of sex-based features even in the presence of psychiatric disease. Previous volumetric work also has strong alignment with our findings, in a single-site study of 5,216 participants, greater volume of the amygdala, caudate, pallidum, putamen, and thalamus was observed in males compared to females, but with females showing greater volume in the accumbens (Ritchie et al., 2018). Furthermore, a meta-analysis of sex differences in brain structure reported larger volumes and higher tissue densities in the bilateral amygdala, hippocampus, and putamen for men (Ruigrok et al., 2014). And finally, in a recent study by Liu and colleagues (2020) involving two large independent samples, they observed consistently greater gray matter volume in men within the amygdala, hippocampus, and putamen.

While both males and females with schizophrenia showed decreased thickness and surface contraction of deep brain regions compared to healthy control participants, the pattern of difference between the male and female contrasts was quite similar and not consistent with our initial hypothesis. In other words, the impact a schizophrenia diagnosis had on deep brain shape did not vary as a function of biological sex. While many researchers have highlighted sex differences in the clinical correlates of schizophrenia (e.g., positive and negative symptoms, cognitive deficits, medication, etc.), few studies have investigated differences in the brain between men and women with the disease (Li et al., 2016; Ochoa et al., 2012; Seeman, 2021). A systematic review of schizophrenia-based fMRI studies by Salehi and colleagues (2024) reported multiple brain-based sex differences in temporal, frontal and hippocampal regions, shedding light on the underlying pathophysiology. However, they speculated that additional work was needed to compare patterns observed in healthy individuals to schizophrenia in order to determine whether disease-related differences were unique. Thus it is possible that brain-based abnormalities in other modalities may also not vary on the basis of biological sex, and perhaps other mechanisms (hormonal, cellular, etc.) could better explain differences in cognition and behavior between men and women with schizophrenia.

Regarding asymmetry, the present study identified exaggerated deep brain asymmetry in schizophrenia men relative to healthy men, which was not observed in schizophrenia women, and was primarily observed in thalamic and hippocampal regions. Furthermore, sex-specific shape asymmetry in healthy individuals (primarily amygdala, caudate and thalamus) was noticeably absent in the male-female schizophrenia comparison, again suggesting the illness perturbs normally expressed variation in brain structure. These findings extend previous work in volumetric asymmetries, such as a large-scale investigation of cortical and subcortical regions by Schijven and colleagues (2023) that noted asymmetric differences between schizophrenia and healthy individuals primarily in several cortical regions, but interestingly did not observe significant sex-specific effects on any asymmetry index. Furthermore, in our previous study with overlapping subjects, Gutman and colleagues (2022) described patterns of exaggerated asymmetry in the hippocampus, amygdala, and thalamus in patients with schizophrenia, although effects were small. In an ENIGMA consortium study of multiple healthy pediatric samples, prominent asymmetries were noted in both males and females with more regions affected in females, but a greater magnitude of asymmetry in males; in both instances the observed effect sizes were small (Kurth et al., 2024). While our work differed with the above literature in some areas, it does suggest that normal sex-based asymmetries in deep brain morphology become attenuated in the context of schizophrenia, thus disrupting a typical neurobiological presentation.

Early studies on structural abnormalities and pathophysiological implications for patients with schizophrenia suggest that brain morphology is impacted by environmental and genetic risk factors, course of illness, and treatment (Buckley, 2005). In the context of the illness, these features interact with proposed mechanisms contributing to sex differences in neuroanatomical organization, including evolutionary, genetic, and environmental influences (DeCasien et al., 2022). More recently, research has focused on the impact of gonadal steroids and sex chromosome dosage on brain organization. Specifically, increases in surface area of the dorsal caudate, dorsal thalamus, and rostro-caudal extremes of the hippocampus, as well as decreases in volumes of the pallidum and amygdala and decreases in cortical thickness, have been associated with carriage of supernumerary X chromosomes compared to karyotypically normal males and females (Arnold, 2020; Nadig et al., 2018). The present study’s findings highlight a pattern of greater thickness/surface contraction across most regions of interest in males compared to females regardless of diagnosis, indicating sex chromosome dosage may be a potent mechanism contributing to some of the observed sex differences, but not provide a comprehensive explanation for differences observed in the caudate, thalamus, and hippocampus. While effect sizes for genetic variants are small, epigenetic research has reported possible interactions with environmental factors, including antenatal maternal virus infections or hypoxia during neurodevelopment, that can contribute to hippocampal morphometry in patients with schizophrenia (Schmitt et al., 2014), a pattern that was consistently observed between males and females in the present study. While sex differences in global and regional brain anatomy are reproducible, researchers are still uncertain of their causal factors or potential connections with human behavior (DeCasien et al., 2022).

The clinical presentation of schizophrenia has noted to differ between men and women with the illness in terms of symptom type/severity, diagnostic timeline, and medication treatment (Giordano et al., 2021; Li et al., 2021; Sommer et al., 2020). Researchers have examined how shared neurobiological patterns can predict different behaviors in men and women. Of particular interest are the sex-specific effects of hippocampus neurons that could potentially contribute to differences in clinical presentation between men and women with depression (Mulvey et al., 2023). These researchers investigated how risk-associated functional variants interact with biological sex and affect men and women differently. Given that hippocampal deformation has also been observed as a biomarker of patients with schizophrenia (Egloff et al., 2018), and that men are more commonly diagnosed with psychotic-spectrum disorders (Li et al., 2016), further research could explore how sex-by-allele effects can underlie sex bias in the disease. Furthermore, Dhamala and colleagues (2023) highlighted the neurobiological correlates of many internalizing and externalizing behaviors that are observed across sexes, although they manifest differently between men and women with psychiatric illness.

The present study did not find statistically meaningful relationships between negative symptoms or duration of illness with deep brain shape for men or women with schizophrenia. However, significant relationships were observed between positive symptoms (SAPS only) and shape deformation in the bilateral caudate, right hippocampus, and right amygdala in both men and women, the pattern of which was highly similar between them suggesting no unique sex-specific signatures. A similar finding was noted in the antipsychotic models where CPZ significantly related with deformation in the caudate, accumbens, hippocampus, amygdala, and thalamus in both men and women, but no differential pattern. Previous literature on antipsychotic use and the brain is largely inconclusive; one study noted associations between antipsychotics and increased volume in the anterior cingulate cortex and right putamen, as well as decreased volume in the frontal and temporal cortices (Chen et al., 2021; Torres et al., 2013). On the other hand, a review by Lawrie (2018) suggested that the effects of antipsychotic drugs vary across patients and do not appear to cause adverse structural changes. When considered in light of this study’s findings, the impact of medication use on deep brain shape does not appear to have a sex-specific feature and may be related more to individual or type/dosage factors.

In sum, this meta-analytic study aimed to understand whether the presence of schizophrenia would result in alterations to standard sex-related morphological differences of deep brain structures critical to various cognitive, emotional, and behavioral processes. The primary finding is that sex differences in brain morphology are robust and fundamental across both healthy and schizophrenia states, suggesting any differences are not artifacts of disease, but inherent characteristics of female and male brains. This observation emphasizes the need to control, or account for, sex as a key variable when studying the neurobiology of schizophrenia. Furthermore, as schizophrenia does not appear to uniquely alter sex-related differences in brain structure it suggests the neuroanatomical impact of the illness is perhaps more uniform across men and women. It may also suggest that various neuroprotective or vulnerability factors are evenly distributed in both sexes, which would shift schizophrenia-focused research towards underlying mechanisms that are not sex dependent. Understanding why schizophrenia does not exacerbate, or alter, typical sex differences patterns in brain morphology could aid insights into the biological underpinnings of the disease, such as if there is a contribution to a standardized pattern of disease progression across sexes.

Given that men and women with schizophrenia present differently from a clinical standpoint, yet the observed brain morphology patterns represent that observed in healthy individuals, treatments and diagnostic criteria might need reconsideration to better account for inherent sex differences that persist irrespective of disease. Recent theories have explored schizophrenia subtypes based on predominant symptomatic experiences. An important avenue for future research, therefore, would be to explore whether sex differences in the brain map onto traits that serve as risk factors for severe mental illness, or even specific patterns of clinical presentation. Furthermore, the lack of research on distal and proximal influences of sex differences in the brain leads to challenges as it pertains to interpreting patterns of shape morphology. Future research could focus on more granular (e.g., molecular and genetic) or macro (e.g., connectivity) bases of deep brain morphology to determine why sex differences persist in the face of schizophrenia and how these differences could relate to resilience or susceptibility to the disease..

The present study had several limitations that should be considered in future work. Given the neuroimaging and clinical data was collected from multiple sites, a meta-analytic approach was used to generate consensus maps. Meta-analysis is limited in that the data may not be homogeneous, and there is a potential for nonlinear correlations. To target these concerns, all subcortical shape measures were processed using the validated ENIGMA-Shape pipeline where a set of standardized scripts were used on data from all contributing sites to compute mass univariate statistics. Furthermore, methodological considerations, in part, accounted for this limitation by aggregating effect sizes across sites. Another approach for a study that reviews data from multiple sites is mega-analysis, which pools raw data across studies. In this sense, site-level effects are modeled as random effects. Meta-analyses do not assume that data from contributing sites have the same mean and variance. Another consideration is the noise that is generated by the FreeSurfer software, which can impact shape measurements that are based on vertex-wise information. This limitation was mostly addressed through quality assurance procedures that were completed by individual raters and the USC Imaging Genetics Center.

## Conclusion

In conclusion, this meta-analysis of deep brain regions revealed patterns of shape deformation that provide greater insights into sex differences in healthy individuals and patients with schizophrenia. Findings revealed significant patterns of deformation between patients with schizophrenia and healthy individuals, as well as significant differences between male and female brains. However, a diagnosis of schizophrenia does not appear to impact the brain differently based on biological sex. These results provide important avenues for future research and can enhance understanding of the unique neurobiological contributions of psychosis as they pertain to biological sex, the application of which can strengthen assessment, improve diagnostic approaches, and allow for more effective personalized treatment of patients with schizophrenia.

## Acknowledgements

Data used in this publication was included in a poster presentation at the 2023 conference for the Schizophrenia International Research Society. Research reported in this publication was supported by the National Institute of Biomedical Imaging and Bioengineering (NIBIB) of the National Institutes of Health under Award Number 5U54EB020403 and 5R01MH116147.

The CIAM study (FMH - PI) was supported by the University Research Committee, University of Cape Town and the National Research Foundation, South Africa.

The Dublin study was supported by grant funding from the Irish Health Research Board (grant number HRA_POR/2012/54) and Science Foundation Ireland (grant numbers 12/IP/1359 and 08/IN.1/B1916).

The FBIRN study was supported by the National Center for Research Resources at the National Institutes of Health (grant numbers: NIH 1 U24 RR021992 (Function Biomedical Informatics Research Network) and NIH 1 U24 RR025736-01 (Biomedical Informatics Research Network Coordinating Center; http://www.birncommunity.org).

The UCISZ study was supported by the National Institutes of Mental Health grant number R21MH097196 to TGMvE. FBIRN and UCISZ data were processed by the UCI High Performance Computing cluster supported by Joseph Farran, Harry Mangalam, and Adam Brenner and the National Center for Research Resources and the National Center for Advancing Translational Sciences, National Institutes of Health, through Grant UL1 TR000153. FBIRN thanks Mrs. Liv McMillan for overall study coordination.

The FIDMAG study was supported by Miguel Servet Research Contract MS14/00041 and Research Project PI14/00292 from the Plan Nacional de I+D+i 2013–2016, the Instituto de Salud Carlos III-Subdirección General de Evaluación y Fomento de la Investigación and the European Regional Development Fund (FEDER), Juan de la Cierva-formación contract (FJCI-2015-25278), and Cibersam.

The Galway study was supported by grant funding from the Health Research Board (grant number HRA_POR/2011/100) and the Wellcome Trust (grant number 072894/2/03/Z).

The Hubin study was supported by the Swedish Research Council (grant numbers K2009-62X-15077-06-3 and K2012-61X-15077-09-3), the Karolinska Institutet and the Knut and Alice Wallenberg Foundation.

The KaSP study was supported by grants from the Swedish Medical Research Council (SE: 2009-7053; 2013-2838; SC: 523-2014-3467), the Swedish Brain Foundation, Åhlén-siftelsen, Svenska Läkaresällskapet, Petrus och Augusta Hedlunds Stiftelse, Torsten Söderbergs Stiftelse, the AstraZeneca-Karolinska Institutet Joint Research Program in Translational Science, Söderbergs Königska Stiftelse, Professor Bror Gadelius Minne, Knut och Alice Wallenbergs stiftelse, Stockholm County Council (ALF and PPG), KID-funding from the Karolinska Institute.

The MCIC study was supported by the National Institutes of Health (NIH/NCRR P41RR14075 and R01EB005846 (to Vince D. Calhoun)), the Department of Energy (DE-FG02-99ER62764), the Mind Research Network, the Morphometry BIRN (1U24, RR021382A), the Function BIRN (U24RR021992-01, NIH.NCRR MO1 RR025758-01, NIMH 1RC1MH089257 to Vince D. Calhoun), the Deutsche Forschungsgemeinschaft (research fellowship to Stefan Ehrlich), and a NARSAD Young Investigator Award (to Stefan Ehrlich).

The NU study was supported by NIH grants P50 MH071616, R01 MH056584, R01 MH084803 (Wang PI), T32 NS047987 (Cobia PI), U01 MH097435 (Wang, Turner, Ambite, Potkin PIs), R01 EB020062 (Miller, Paulsen, Mostfosky, Wang PIs), NSF 1636893 (Pestilli, Wang, Saykin, Sporns PIs), NSF 1734853 (Pestilli, Garyfallidis, Henschel, Wang, Dinov PIs).

The Olin study was supported by R37MH43375 and R01MH074797.

The PAFIP study was supported by Instituto de Salud Carlos III, FIS 00/3095, 01/3129, PI020499, PI060507, PI10/00183, the SENY Fundació Research Grant CI 2005-0308007, and the Fundación Marqués de Valdecilla API07/011.

The TOP study was supported by the Research Council of Norway (#213837, #217776, #223273), the South-East Norway Health Authority (2013-123), and the KG Jebsen Foundation.

The UPenn study was supported by National Institute of Mental Health grants MH064045, MH 60722, MH019112, and MH085096 (DHW). Theodore D. Satterthwaite was supported by MH098130 and by the Marc Rapport Family through NARSAD.

The SLF Rome study was supported by the Italian Ministry of Health grant RC-12-13-14-15-16-17-18/A.

The NARSAD and Wellcome studies were supported by grants from FAPESP-Brazil (#2009/14891-9, 2010/18672-7, 2012/23796-2 & 2013/03905-4), CNPq-Brazil (#478466/2009 & 480370/2009), the Wellcome Trust (UK) and the Brain & Behavior Research Foundation (2010 NARSAD Independent Investigator Award granted to Geraldo F. Busatto).

The Queensland Twin IMaging (QTIM) study was supported by the National Institutes of Health (R01 HD HD050735, 1U54EB020403-01, subaward no. 56929223) and the National Health and Medical Research Council (1009064, 496682).

Research reported in this publication was also supported by the following VA grants (VA I01 CX000497 and Senior Research Career Scientist to JMF) and National Institutes of Health grants: U54 EB020403 to PMT, R01 MH116147, U24 RR21992, R21 MH097196, and TR000153 to TGMvE, S10 OD023696 and R01EB015611 to PK, T32 AG058507 and 5T32 MH073526 to CRKC, R01 MH117601 to NJ, R01 DA053028 to LW and JT. The content is solely the responsibility of the authors and does not necessarily represent the official views of the funding agencies.

